# Complex interactions between weather, and microbial and physiochemical water quality impact the likelihood of detecting foodborne pathogens in agricultural water

**DOI:** 10.1101/2020.01.02.892851

**Authors:** Daniel Weller, Natalie Brassill, Channah Rock, Renata Ivanek, Erika Mudrak, Sherry Roof, Erika Ganda, Martin Wiedmann

**Author notes:** Address correspondence to Martin Wiedmann,. Daniel Weller, Department of Biostatistics, University of Rochester, Rochester, NY. Erika Ganda, Department of Animal Science, Pennsylvania State University, State College, PA.

## Abstract

Agricultural water is an important source of foodborne pathogens on produce farms. Managing water-associated risks does not lend itself to one-size-fits-all approaches due to the heterogeneous nature of freshwater environments, and because environmental conditions affect the likelihood of pathogen contamination and the relationship between indicator organism levels (e.g., *E. coli*) and pathogen presence. To improve our ability to develop location-specific risk management practices, a study was conducted in two produce-growing regions to (i) characterize the relationship between *E. coli* levels and pathogen presence in agricultural water, and (ii) identify environmental factors associated with pathogen detection. Three AZ and six NY waterways were sampled longitudinally using 10-L grab samples (GS) and 24-h Moore swabs (MS). Regression showed that the likelihood of *Salmonella* detection (Odds Ratio [OR]=2.18), and *eaeA-stx* codetection (OR=6.49) was significantly greater for MS compared to GS, while the likelihood of detecting *L. monocytogenes* was not. Regression also showed that *eaeA-stx* codetection in AZ (OR=50.2) and NY (OR=18.4), and *Salmonella* detection in AZ (OR=4.4) were significantly associated with *E. coli* levels, while *Salmonella* detection in NY was not. Random forest analysis indicated that interactions between environmental factors (e.g., rainfall, temperature, turbidity) (i) were associated with likelihood of pathogen detection and (ii) mediated the relationship between *E. coli* levels and likelihood of pathogen detection. Our findings suggest that (i) environmental heterogeneity, including interactions between factors, affects microbial water quality, and (ii) *E. coli* levels alone may not be a suitable indicator of the food safety risks. Instead, targeted methods that utilize environmental and microbial data (e.g., models that use turbidity and *E. coli* levels to predict when there is a high or low risk of surface water being contaminated by pathogens) are needed to assess and mitigate the food safety risks associated with preharvest water use. By identifying environmental factors associated with an increased likelihood of detecting pathogens in agricultural water, this study provides information that (i) can be used to assess when pathogen contamination of agricultural water is likely to occur, and (ii) facilitate development of targeted interventions for individual water sources, providing an alternative to existing one-size-fits-all approaches.

---

Preharvest surface water use for produce production (e.g., irrigation, fertigation, pesticide application, dust abatement) has repeatedly been identified as a factor associated with an increased likelihood of foodborne pathogen contamination of fresh produce [e.g., (Holvoet et al., 2014; Mody et al., 2011; Strawn et al., 2013b; Weller et al., 2015b)]. This is largely because (i) surface water can act as a source (McEgan et al., 2013a; Micallef et al., 2012) and transmission pathway (Girardin et al., 2005; Mody et al., 2011; Weller et al., 2015b) for foodborne pathogens in farm environments, and (ii) the use of pathogen-contaminated water can transfer pathogens to produce directly (Erickson et al., 2010; Guan et al., 2001) and indirectly [e.g., through contamination of the farm environment (Ibenyassine et al., 2006; Oliveira et al., 2012)]. In fact, irrigation with untreated surface water has been repeatedly associated with the isolation of foodborne pathogens, such as *Listeria monocytogenes*, from preharvest environments (Guan et al., 2001; Holvoet et al., 2014; Strawn et al., 2013b; Weller et al., 2015b), and identified as a potential cause of outbreaks linked to produce (Baloch, 2014; Centers for Disease Control and Prevention (U.S.), 2018; Food and Drug Administration (U.S.), 2018; Mody et al., 2011). Thus, mitigating the food safety risks associated with preharvest surface water use is a priority.

Recognizing the role of surface water as a source of foodborne pathogens, the US Food and Drug Administration proposed microbial water quality standards as part of the Food Safety Modernization Act’s (FSMA) Produce Safety Rule. The proposed standard states that *E. coli* levels in surface water directly applied to preharvest produce cannot exceed a geometric mean of 126 CFU/100-mL and a statistical threshold value (STV) of 410 CFU/100-mL; these values are calculated using 20 samples collected over a 2 to 4 year period. However, understanding and complying with the proposed standard while ensuring water availability has been cited in industry magazines and grower surveys as a challenge facing growers (Alexander, 2015; Dery et al., 2019; Wall et al., 2019). For example, interpretation of *E. coli* test results is complicated by temporal variation in microbial water quality (Goyal et al., 1977; Hipsey et al., 2008; Pandey et al., 2012; Payment and Locas). Since the samples used to determine if a water source meets the proposed standard can be collected up to 4 years before the water source is used for produce production, meeting the standard also may be a poor approximation of water quality at time of use (Havelaar et al., 2017; Truitt et al., 2018). Acceptance of *E. coli*-based water standards is further complicated by conflicting data on the relationship between *E. coli* levels and pathogen presence in the literature (Benjamin et al., 2013; Economou et al., 2013; Harwood et al., 2005; McEgan et al., 2013a; Pachepsky et al., 2015; Wilkes et al., 2009). While some studies argue that high *E. coli* levels are associated with an increased likelihood of detecting pathogens in agricultural water (Edberg et al., 2000; Payment and Locas; Wilkes et al., 2009), other studies disagree (Antaki et al., 2016; Benjamin et al., 2013; Harwood et al., 2005; Pachepsky et al., 2015). While this differentiation is often not made, these observations are consistent with the fact that *E. coli* is considered an indicator of potential fecal contamination, and not an “index organism” [detection of an index organism suggests the presence of an ecologically similar or closely-related pathogen (Busta et al., 2006; Chapin et al., 2014)]. Despite this, it is important to understand the relationship between *E. coli* levels and foodborne pathogen contamination of agricultural water since data on *E. coli* levels are used to guide efforts to mitigate the microbial food safety risks associated with preharvest surface water use (e.g., to decide if and when corrective measures such as water treatment should be implemented). Thus, data in the relationship between *E. coli* levels and foodborne pathogen contamination of agricultural water are essential for identifying when and where *E. coli* levels can be used (alone or in conjunction with other data) to manage the food safety risks associated with preharvest water use. Promising alternative approaches include models that predict risk of pathogen presence at specific times and sampling locations along a waterway by using a variety of different, spatially explicit input data. However, to develop these alternative approaches additional data on factors (e.g., weather, physiochemical water quality) that drive temporal variation in *E. coli* levels and pathogen presence in different regions and under different conditions is needed.

Like *E. coli* levels, the prevalence of key foodborne pathogens in surface water also varies between studies and over time. For example, 30% and 63% of surface water samples collected from New York produce farms in 2010 (Weller et al., 2015b) and 2015 (Weller et al., 2015a), respectively, were *L. monocytogenes-*positive. Similarly, 29% (Kayed, 2004), 58% (Ijabadeniyi et al., 2011), and 67% (Castillo et al., 2004), of water samples collected from canals in Arizona, South Africa and Texas, respectively, were *Salmonella*-positive. While this variability may be due to the spatiotemporal heterogeneity of farm and freshwater environments (Benjamin et al., 2013; McEgan et al., 2013a), the methods used to collect and process water samples may also affect reported pathogen prevalence (Benjamin et al., 2013; Colburn et al., 1990; Hoganson and Elliott, 1972). Thus, understanding how temporal variation in environmental factors affects microbial water quality, and how sampling methods affect our ability to detect pathogens in agricultural water is essential to effectively manage the food safety risks associated with preharvest water use. Thus, the objectives of our study were to: (i) quantitatively assess the association between *E. coli* levels and pathogen presence in surface water sources used for produce production; (ii) identify and rank environmental factors associated with detecting pathogens in these waterways; and (iii) compare the ability of two sampling methods, 24-h Moore swabs (MS) and 10-L grab samples (GS), to detect pathogens in agricultural water. Since environmental conditions are highly variable between regions, multi-region studies are needed to ensure that findings are translatable to outside the study area, and to allow researchers to identify region-specific and consistent risk factors. Thus, two produce-growing regions, southwestern Arizona and western New York, were sampled as part of the study reported here.

## MATERIALS AND METHODS

### Study design

A longitudinal study was conducted in AZ and NY; sampling in AZ occurred between February and December 2017 while sampling in NY occurred between May and September 2017; these time frames correspond to the growing season in each region. At each sampling, we collected a set of Moore swabs [MS] and a set of grab samples [GS]. Each GS set consisted of three 10-L GS [one 10-L GS for detection of each of the three targets (*Listeria, Salmonella*, and the *stx*/*eaeA* genes) and one 1-L GS for enumeration of *E. coli* levels. GS were collected from the middle of each channel and approx. 15 cm (6 inches) below the water surface. Each MS set consisted of three swabs (one swab for detection of each of the target groups). Gloves (Nasco, Fort Atkinson, WI) were changed for each sample collected, and sampling materials were sprayed with 70% ethanol in between all sample collections. All samples were transported on ice, and stored at 4°C until processing. The 10-L GS used for pathogen detection were processed within 18 h of collection, while the 1-L GS used for *E. coli* enumeration were processed within 6 h of collection asper manufactures’ instructions.

Samples were collected using a “1-week sampling” scheme and a “1-day sampling” scheme to maximize temporal coverage (Fig S1). On the first day of each 1-week sampling, a MS set was anchored in the waterway and a GS set was collected. When the first MS set was collected 24 h later, a second GS set was collected and a second MS set was deployed. This was repeated daily for up to 6 d. During 1-day sampling, a MS set was placed in the waterway for 24 h. During this 24 h period, six GS sets were collected between 6 am and 8 pm approx. 2.5 hours apart. One-week sampling was performed on eight waterways (2 AZ canals and 6 NY streams) three times each, while 1-day sampling was performed on seven waterways (2 AZ canals and 5 NY streams) three times each. One-day sampling was performed on fewer waterways than the 1-week sampling due to the substantial time needed to perform a single 1-day sampling. While 1-day sampling was performed once on a third AZ canal, this canal was removed from the study for logistical reasons after the first 1-day sampling.

### Waterway Enrollment and Spatial Data Acquisition

Watershed delineation and all other spatial analyses were performed in ArcGIS version 10.2. Remotely sensed data (e.g., flow accumulation rasters) were obtained from publicly available databases to facilitate waterway enrollment. Hydrological, land use, road, and other spatial data were downloaded from federal and state geodata portals (e.g., www.usgs.gov/core-science-systems/ngp/national-hydrography; cugir.library.cornell.edu/).

Sampling sites in NY were enrolled by randomly selecting six streams with non-overlapping watersheds from all eligible streams in the study region. Specifically, streams were enrolled by identifying watersheds with an area of ≥15 km^2^ and where produce was grown in ≥ 4 of the last 8 years based on USDA Cropscape data (Boryan et al., 2011; Han et al., 2012, 2014). We then randomly selected 6 publicly accessible locations that were ≤400 m from a produce field along streams in these watersheds (Fig S2). Publicly accessible sites were locations near stream-road intersections, on public land (e.g., parks, Cornell farms), or with public-right-of-way (e.g., fishing access). Sampling sites in AZ were enrolled to represent the diversity of canal types in produce-growing regions of Arizona and based on the willingness of the irrigation district to provide access to the study site. While sites were selected so that they were < 400 m from an active produce farms, sites were not selected using other environmental or geographic criteria. Since access to the canals was dependent on buy-in from the irrigation districts, random site selection could not be performed in AZ, which may have resulted in selection bias.

### Metadata Collection

Every time a sample was collected, metadata were also collected. Specifically, data on dissolved oxygen levels, pH, conductivity, and water temperature were measured in-field using a Hach HQ40d meter (Loveland, CO); turbidity was measured in the laboratory using a turbidimeter (Hach). Flow rate in NY was measured 6 inches below the water surface using a flow meter (Global Water Instrumentation Inc., Cordova, CA), while surface flow was estimated in AZ using the float method described in Gore (Gore, 2006). Meteorological data were obtained from the weather station (cals.arizona.edu/AZMET; www.nysmesonet.org; newa.cornell.edu) closest to each site; the mean distance of the stations to the sites was 8.9 km (range = 0.4 to 25.5 km). Data were downloaded for the entire growing season in each state. Avg., min., and max. air temperature, avg. relative humidity, avg. solar radiation, and avg. wind speed were calculated for 0 to 1, 0 to 2, 0 to 3, 0 to 4, and 0 to 5 days before sample collection (BSC). Total rainfall was calculated using non-overlapping time periods (i.e., 0 to 1, 1 to 2, 2 to 3, 3 to 4, and 4 to 5 d BSC). Since no rain fell in Arizona during the time periods considered, rainfall factors were only included in downstream analyses when *E. coli* levels or pathogen detection in New York was the outcome.

### Grab Sample [GS] Processing

The three 10-L GS were filtered using modified Moore swabs (mMS) as previously described (Sbodio et al., 2013); however, unlike previous studies that used a peristaltic pump to move water through the mMS cassette, we used a gravity-based system. After all 10-L of water were filtered, the mMS was transferred to a Whirl-Pak bag and processed as described below. A 100-mL aliquot of the 1-L GS was used for *E. coli* enumeration, which was performed using the Colilert Quanti-Tray 2000 kit (IDEXX, Westbrook, ME) per manufacturer instructions.

### *Listeria* Enrichment and Isolation

*Listeria* enrichment and isolation were performed as previously described (Weller et al., 2015a). Briefly, 225 mL of buffered *Listeria* enrichment broth (Becton Dickinson, Franklin Lakes, NJ) were added to each Whirl-pak containing a MS or mMS. Following incubation at 30°C for 4 h, *Listeria* selective enrichment supplement (Oxoid, Cambridge, UK) was added to each enrichment. After incubating at 30°C for a total of 24 h and 48 h, 50 µl of enrichment were streaked onto *L. monocytogenes* plating medium (LMPM; Biosynth International, Itasca, IL) and Modified Oxford agar (MOX; Becton Dickinson), which were incubated at 35°C and 30°C, respectively, for 48 h. Following incubation, up to 4 presumptive *Listeria* colonies were sub-streaked from MOX to LMPM and incubated at 35°C for 48 h. From all LMPM plates, up to 2 presumptive *Listeria* (excluding *L. monocytogenes*) colonies and up to 2 presumptive *L. monocytogenes* colonies were sub-streaked onto brain-heart infusion plates (BHI; Becton Dickinson). Fewer than the maximum number of colonies were selected if sufficient colonies were not available for a given sample. The BHI plates were incubated at 37°C for 24 h. The species of one presumptive *Listeria* (excluding *L. monocytogenes*) colony and one presumptive *L. monocytogenes* colony per sample was determined by PCR amplification and sequencing of the partial *sigB* gene (Bundrant et al., 2011; Den Bakker et al., 2010; Nightingale et al., 2005). Positive [FSL R3-0001 (Roberts and Wiedmann, 2006)] and negative (uninoculated media) controls were processed in parallel with the samples. All isolates were preserved at −80°C.

### *Salmonella* Enrichment and Isolation

Two-hundred and twenty-five mL of buffered peptone water supplemented with novobiocin (final concentration of 20 mg/L; BPW+N) was added to each Whirl-pak containing a MS or mMS. Following incubation at 35°C for 24 h, a *Salmonella* PCR-screen was performed using a real-time BAX *Salmonella* assay (Hygiena, Wilmington, DE). BAX negative samples were considered *Salmonella*-negative, while BAX positive samples were culture-confirmed as *Salmonella*-positive as previously described (Strawn et al., 2013a). Briefly, 1 mL of the BPW+N enrichment was added to 9 mL of tetrathionate broth (TT; Oxoid) supplemented with 200 uL of I2-KI and 100 uL of Brilliant Green. In parallel, 0.1 mL of the BPW+N enrichment was added to 9.9 mL of Rappaport Vassiliadis broth (RV; Acros Organic, Geel, Belgium). After incubating the TT and RV broth in a shaking water at 42°C bath for 24 h, 50 uL of each broth were streaked separately onto *Salmonella* CHROMagar (DRG International, Springfield, NJ) and xylose lysine deoxycholate agar (XLD; Neogen, Lansing, MI) plates. The CHROMagar and XLD plates were incubated for 24 h at 37°C and 35°C, respectively. Following incubation, up to 12 presumptive *Salmonella* colonies per sample were confirmed as *Salmonella* by PCR amplification of the *invA* gene (Kim et al., 2007). Specifically, four presumptive *Salmonella* colonies (mauve colonies on CHROMagar or black colonies on XLD) were selected for PCR-confirmation; if possible, two colonies per media were selected. If there were no presumptive positive colonies on the CHROMagar or XLD plates then up to 12 blue colonies on CHROMagar and/or red colonies on XLD were selected for PCR-confirmation. Positive [media inoculated with *Salmonella* Typhimurium (FSL F6-0826)] and negative (uninoculated media) controls were processed in parallel with field samples. All isolates were preserved at −80°C.

### *eaeA* and *stx* Codetection

A PCR-screen for the *eaeA* and *stx* genes was performed using a real-time BAX Shiga-toxin producing *E. coli* (STEC) assay (Hygiena); this is a proprietary assay that detects the presence of the *stx1* and *stx2* genes. Sample enrichment and processing were performed per manufacturer’s instructions. Briefly, 250 mL of tryptic soy broth supplemented with casamino acids and novobiocin to a final concentration of 10 g/L and 8 mg/L, respectively, (TSB+N) was added to each Whirl-pak. Following incubation at 41°C for 24 h, the BAX assay was performed per the manufacturer’s instructions.

### Statistical Analyses

All analyses were performed in R (version 3.4.2; R Core Team, Vienna, Austria). Changes in environmental conditions over the course of the study were visualized by plotting each factor over time. Correlation between environmental factors was quantified and visualized as previously described (Wei, 2013; Weller et al., 2015a). The prevalence of each of the target organisms [*Listeria* spp. (including *L. monocytogenes*), *L. monocytogenes*, and *Salmonella*] as well as the prevalence of *eaeA* and *stx* codetectiondo not was determined. The geometric mean of *E. coli* (MPN/100-mL) was calculated for each of the sampled waterways and for each state. General linear mixed modeling followed by Tukey’s HSD was used to compare *E. coli* levels between waterways using the lme4 (Bates et al., 2014) and emmeans (Lenth, 2018) packages. The outcome of the model was the log_10_ MPN of *E. coli*/100-mL, the fixed effect was site, and the random effects were year-day and state. Year-day is the number of days since Jan. 1^st^ (e.g., Jan. 1^st^ has is year-day 0, Jan. 2^nd^ is year-day 1).

#### Comparison of Pathogen Detection by 24-h MS and Paired 10-L GS

In this study, we used two sampling methods (24-h MS and 10-L GS). Each MS collected as part of 1-week sampling had between 1 and 2 paired GS, while MS collected as part of the 1-day sampling had between 6 and 7 paired GS (Fig S1). A sampling day was defined as the 24-h the MS was in the stream. For each sampling day, we determined if the MS and/or one of its paired GS detected a given target. Separately, we used generalized linear mixed models to determine if MS were significantly more or less likely to detect a given target compared to a single paired GS (Bates et al., 2014). Since the outcome of the mixed models was binary we used a binomial distribution with a logit link. The explanatory variable was sample type (GS was the reference level). Site nested in state, and year-day were included as random effects. Since the ability of a MS compared to a paired GS to detect pathogen contamination in a given waterway at a given time should not differ between states, AZ and NY data were combined for these analyses.

#### Random Forest Analysis

Random forest analysis was performed separately to identify and rank factors associated with *Salmonella, Listeria* spp. and *L. monocytogenes* isolation, and *eaeA-stx* codetection in each sample type. Random forest analysis was also performed to identify and rank factors associated with *E. coli* levels in GS. Random forest as opposed to other methods (e.g., regression analysis) was chosen as random forests rank factors based on the strength of their association with the outcome but do not generate effect estimates or odds ratios to quantify the strength of these associations. This is important since, as our study shows, there is substantial variability in water quality within a waterway over time, and as such the time span of our study (one growing season) was insufficient to generate reliable effect estimates. Moreover, repeated, three-fold cross-validation was used during random forest development to reduce overfitting and to give insights into how well our findings generalize to independent datasets. AZ and NY data were analyzed separately to allow for identification of region-specific factors that were associated with pathogen detection and *E. coli* levels. For each RF, environmental factors (see Tables S1-2 for a complete list) were included as explanatory factors. Year-day and sample site ID were also included as a proxy for unmeasured spatiotemporal factors. Since this is a hypothesis-generating study, five overlapping periods (0-1, 0-2, 0-3, 0-4, and 0-5 d before sample collection) were used to calculate the values for the weather factors with the exception of rainfall; separate random forests were then run for each combination of time period, outcome, state, and sample type.

Unbiased conditional random forest analysis was performed using the party package and controls recommended by the package authors (Boulesteix et al., 2015; Strobl et al., 2007b, 2007a, 2009). For each random forest, repeated 10-fold cross-validation was performed to tune hyperparameters and to calculate either the Kappa score (Kuhn) for forests where the outcome was categorical, or the coefficient of determination (R^2^) for forests where the outcome was continuous. The random forest with the highest Kappa score for each combination of outcome, state, and sample type is discussed in-text. Factor rankings for all random forests are reported in Tables S3-6, and the variable importance scores for all random forests are available at github.com/wellerd2/PAWQ-2017. For random forests where the outcome was binary and imbalanced (prevalence of positive samples was <40% or >60%) upsampling was performed (Kuhn). Random forest results were interpreted by quantifying conditional variable importance (VI); conditional VI was calculated because multiple explanatory factors were correlated [Fig. S3-S4; (Strobl et al., 2008, 2009)]. A higher VI, relative to all other factors in the random forest, indicated a stronger association between outcome and factor. Variables with VI ≤0 were not associated with the outcome. Since VI is relative, normalized variable importance measures (NVI) were calculated to facilitate interpretation and visualization of the results. For each combination of outcome, state, and sample type the random forest with the highest Kappa score was identified and partial dependence plots (PDPs) were developed to graphically characterize (i) the relationships between top-ranked factors and the outcome, and (ii) the impact of two-way interactions between factors on the outcome (Greenwell, 2017). Interactions were defined as occurring if the marginal effect of one factor on the outcome was not constant over all values of a second factor (Boulesteix et al., 2015). Due to the observational nature of the study reported here, caution should be exercised when interpreting the PDPs. For example, some PDPs indicate a polynomial relationship between a factor and an outcome. However, this relationship may be (i) due to the existence of an optimal range for the target to contaminate, survive or be detected in surface water, (ii) due to the impact of an unmeasured confounder, or (iii) an artifact of sampling and the observational nature of the study. Thus, determining the exact relationship between factors and outcomes is outside the scope of this study; however, all available data on potentially confounding factors were included in the random forests in an attempt to control for this limitation.

#### Characterizing the relationship between E. coli levels and pathogen detection

Generalized linear mixed models (Bates et al., 2014) were developed to characterize the relationship between *E. coli* levels and (i) culture-based *Salmonella*, *Listeria* spp., and *L. monocytogenes* isolation from a sample, and (ii) PCR-based codetection of the *eaeA* and *stx* genes in a sample. Since the outcome of the models was binary, we used a binomial distribution with a logit link. The log_10_ MPN of *E. coli*/100-mL was included as a fixed effect, while year-day and site were included as random effects. Separately, bootstrapping was used to simulate water sampling and create a microbial water quality profile (MWQP) composed of 20 samples (N=10,000 MWQPs per waterway). The simulated MWQPs were then used to quantify the ability of the proposed FSMA standard [geometric mean <126 CFUs/100-mL and STV <410 CFUs/100-mL; (Food and Drug Administration, 2015)] to identify waterways with a high or low risk of pathogen presence at time of water use. The last GS selected for inclusion in each subset (the 20^th^ sample selected) represented microbial water quality at the time of water use (e.g., if the 20^th^ GS selected was *Salmonella*-positive then the water source was considered *Salmonella*-positive at time of water use). The sensitivity, specificity, and diagnostic odds ratio (DOR) were calculated to characterize the predictive accuracy of the proposed standard for each target. AZ and NY data were analyzed separately since differences in environmental conditions and water type (managed canals versus free-flowing streams) may affect the relationship between pathogen detection and *E. coli* levels.

### Data Availability

The R code and output from the random forests are available at https://github.com/wellerd2/PAWQ-2017. The raw data is available upon request with some restrictions (e.g., location of sampling sites cannot be released); data requests should be directed to Martin Wiedmann or Daniel Weller.

## RESULTS

In total, 1,053 grab samples (GS) were collected and analyzed as part of our study [257 10-L GS for *Listeria* isolation, 258 10-L GS for *Salmonella* isolation, 264 10-L GS for *eaeA* and *stx* detection, and 264 1-L GS for enumeration of *E. coli* levels (Table 1)]. Additionally, 362 MS were collected and analyzed for pathogen presence [120 for *Listeria* isolation, 121 for *Salmonella* isolation, and 121 for *eaeA-stx* codetection (Table 2)]. Different numbers of samples were analyzed for different targets due to the loss of samples in the field (e.g., some MS were lost during storms and to human tampering, some containers used for collection of the 10-L GS burst during transport from the field to the lab; some sample sets were removed due to failed control reactions). As a result, we have data on *eaeA-stx* codetection for 121 sampling days, and on *Listeria* and *Salmonella* isolation for 120 sampling days (Table 3); a sampling day is defined as the 24-h period during which a MS was deployed. Figs S3-S6 show correlation between and variation in environmental conditions over the course of the study.

**Table 1:** Summary of grab sample (GS) results.

| Water Source | Prevalence (No. of Positive GS/Total No. of GS) |  |  |  |  | Geometric Mean MPN of *E. coli*/100 mL (Range)<sup>c d</sup> |
| --- | --- | --- | --- | --- | --- | --- |
|  | Culture-Confirmed |  |  | PCR-Screen Positive<sup>b</sup> |  |  |
|  | *L. monocytogenes* | *Listeria* spp.<sup>a</sup> | *Salmonella* | *eaeA* | *stx* |  |
| Arizona |  |  |  |  |  |  |
| Canal A | 3% (1/40) | 3% (1/40) | 40% (16/40) | 92% (33/36) | 58% (21/36) | 217.5 (26.6 - 770.1)<br>DEFG |
| Canal B | 6% (2/36) | 6% (2/36) | 27% (10/37) | 76% (31/41) | 63% (26/41) | 9.6 (1.0 - 47.4)<sup>ABC</sup> |
| Canal C | - | - | - | 83% (5/6) | 0% (0/6) | 4.3 (1.0 - 16.1)<sup>ABC</sup> |
| AZ Total | 4% (3/76) | 4% (3/76) | 34% (26/77) | 83% (69/83) | 57% (47/83) | 38.2 (1.0 - 770.1) |
| New York |  |  |  |  |  |  |
| Stream A | 15% (5/34) | 21% (7/34) | 59% (20/34) | 100% (34/34) | 91% (31/34) | 419.8 (57.6 - >2,419.6)<br>G |
| Stream B | 13% (4/32) | 50% (16/32) | 56% (18/32) | 91% (29/32) | 38% (12/32) | 91.6 (18.5 - 1,413.6)<sup>A</sup><br>D |
| Stream C | 6% (2/33) | 27% (9/33) | 33% (11/33) | 88% (29/33) | 64% (21/33) | 207.6 (23.1 - >2,419.6)<br>C FG |
| Stream D | 21% (7/34) | 50% (17/34) | 41% (14/34) | 100% (34/34) | 76% (26/34) | 221.0 (35.9 - 1,986.3)<br>BC EF |
| Stream E | 21% (7/33) | 85% (28/33) | 33% (11/33) | 94% (31/33) | 70% (23/33) | 108.0 (27.5 - >2,419.6)<br>AB DE |
| Stream F | 13% (2/15) | 53% (8/15) | 40% (6/15) | 93% (14/15) | 80% (12/15) | 175.7 (72.7 - 1,732.9)<br>ABCDEFG |
| NY Total | 15% (27/181) | 47% (85/181) | 44% (80/181) | 94% (171/181) | 69% (125/181) | 181.5 (18.5 - >2,419.6) |
| Total | 12% (30/257) | 34% (88/257) | 41% (106/258) | 91% (240/264) | 65% (172/264) | 109.4 (1.0 - >2,419.6) |

<sup>a</sup> *Listeria* spp. includes *L. monocytogenes*. All *Listeria* isolates from AZ were *L. monocytogenes*. In NY, we isolated *L. booriae*, *L.* *innocua*, *L. marthii*, *L. seeligeri*, and *L. welshimeri* from 2, 9, 10, 28, and 19 NY GS, respectively. We also identified one isolate that did not group with any previously reported *Listeria* species based on sequencing of the *sigB* gene. Eleven GS were positive for both *L.* *monocytogenes* and one other *Listeria* species. <sup>b</sup> In total, 43 AZ GS (i.e., 4 *stx* positive GS were *eaeA* negative) and 125 NY GS (all *stx* positive GS were *eaeA* positive) were positive for *eaeA* and *stx*.
<sup>c</sup> All GS had detectable levels of *E. coli* ( $\geq 1$ MPN/100-mL). Nine NY GS had *E. coli* levels above the detection limit (2419.6 MPN/100 mL). For these GS, a value of 2,500 MPN/100-mL was substituted when estimating the geometric mean. <sup>d</sup> Sites with the same superscript capital letters did not have significantly different *E. coli* levels according to Tukey's HSD.

**Table 2:** Summary of MS (MS) results.

| Water Source | Prevalence (No. of Positive MS/Total No. of MS) |  |  |  |  |
| --- | --- | --- | --- | --- | --- |
|  | Culture-Confirmed |  |  | PCR-Screen Positive<sup>b</sup> |  |
|  | *L. monocytogenes* | *Listeria* spp.<sup>a</sup> | *Salmonella* | *eaeA* | *stx* |
| Arizona |  |  |  |  |  |
| Canal A | 0% (0/17) | 0% (0/17) | 75% (12/16) | 100% (15/15) | 100% (15/15) |
| Canal B | 0% (0/16) | 0% (0/16) | 56% (9/16) | 94% (16/17) | 88% (15/17) |
| Canal C | 0% (0/1) | 0% (0/1) | 0% (0/1) | 100% (1/1) | 0% (0/1) |
| AZ Total | 0% (0/34) | 0% (0/34) | 64% (21/33) | 97% (32/33) | 91% (30/33) |
| New York |  |  |  |  |  |
| Stream A | 6% (1/15) | 19% (3/16) | 56% (9/16) | 100% (16/16) | 100% (16/16) |
| Stream B | 0% (0/15) | 20% (3/15) | 47% (7/15) | 87% (13/15) | 67% (10/15) |
| Stream C | 20% (3/15) | 20% (3/15) | 67% (10/15) | 100% (15/15) | 80% (12/15) |
| Stream D | 0% (0/15) | 13% (2/16) | 63% (10/16) | 100% (16/16) | 94% (15/16) |
| Stream E | 0% (0/14) | 57% (8/14) | 50% (7/14) | 100% (14/14) | 93% (13/14) |
| Stream F | 17% (2/12) | 33% (4/12) | 58% (7/12) | 92% (11/12) | 92% (11/12) |
| NY Total | 7% (6/86) | 27% (23/86) | 57% (50/88) | 97% (85/88) | 88% (77/88) |
| Total | 5% (6/120) | 19% (23/120) | 59% (71/121) | 97% (117/121) | 88% (107/121) |

<sup>a</sup> *Listeria* spp. includes *L. monocytogenes*. In total, we isolated *L. innocua*, *L. marthii*, *L. seeligeri*, and *L. welshimeri* from 2, 2, 11, and 8 NY MS, respectively. All 6 *L. monocytogenes*-positive MS were also positive for one other *Listeria* species.
<sup>b</sup> No MS were positive for *stx* and negative for *eaeA*.

**Table 3:** Comparison of the ability of 24-h Moore swabs (MS) and paired 10-L grab samples (GS) 1368 to detect foodborne pathogens in surface water.

| Target | No. of Detection Events/Total No. of Sampling Days with Paired GS-MS <sup>a</sup> | No. of Events Detected By |  | Disagreement <sup>c</sup> | Regression Results <sup>d</sup> |  |  |  |  |  |
| --- | --- | --- | --- | --- | --- | --- | --- | --- | --- | --- |
|  |  | GS Only <sup>b</sup> | MS Only |  | Fixed Effects |  |  | Variance of Random Effects (SD<sup>g</sup>) |  |  |
|  |  |  |  |  | OR <sup>e</sup> | 95% CI <sup>f</sup> | *P*-value | Year-day | State | Site |
| *L. monocytogenes* | 29/117 | 23 | 5 | 97% (28/29) | 0.39 | 0.16, 0.97 | 0.043 | 0.1 (0.4) | 0.5 (0.7) | 0.0 (0.0) |
| *Listeria* spp.<sup>h</sup> | 61/117 | 38 | 7 | 74% (45/61) | 0.24 | 0.12, 0.48 | <0.001 | 1.4 (1.2) | 3.1 (1.8) | 0.9 (1.0) |
| *Salmonella* | 90/119 | 21 | 30 | 57% (51/90) | 2.18 | 1.37, 3.46 | 0.001 | 0.4 (0.6) | 0.0 (0.0) | 0.0 (0.0) |
| *eaeA* and *stx* | 114/121 | 7 | 23 | 26% (30/114) | 6.49 | 3.10, 13.61 | <0.001 | 1.7 (1.3) | 0.0 (0.0) | 0.6 (0.8) |

<sup>a</sup> A detection event was defined as occurring if the MS or one or more of paired GS tested positive for the target organism. While we collected and tested GS and MS for *Listeria* on 120 sampling days we had paired MS-GS data for 117 samplings days. Similarly, we collected and tested GS and MS for *Salmonella* on 121 sampling days but had paired MS-GS data for 119 samplings days.
<sup>b</sup> GS collected during the 24 h that the MS was in the waterway; each MS had between 1 and 7 paired GS.
<sup>c</sup> Disagreement = (No. of events detected by GS Only + No. of events detected by MS Only)/Total No. of Detection Events
<sup>d</sup> Results of generalized linear mixed models that compared target detection by 24-h MS and individual paired 10-L GS; GS were the reference level.
<sup>e</sup> Odds ratio: Odds of detecting the target in a MS/Odds of detecting the target in a single paired GS
<sup>f</sup> 95% Confidence Interval
<sup>g</sup> Standard Deviation
<sup>h</sup> Includes *L. monocytogenes*

### *E. coli* levels in AZ and NY

Geometric mean *E. coli* levels ranged between 4.3 and 217.5 MPN/100-mL in AZ canals, and between 91.6 and 419.8 MPN/100-mL in NY streams (Table 1). Based on regression analysis, *E. coli* levels varied significantly between waterways (Table 1). For instance, *E. coli* levels in Canal A in AZ were, on average, significantly higher than *E. coli* levels in Canals B and C (Table 1).

For random forests where *E. coli* levels in AZ and NY were the outcome, the random forests with the highest coefficient of determination was based on weather 0-5 days before sample collection (BSC; AZ R^2^ = 0.72; NY R^2^ =0.45; Table S3). For AZ canals, the top-ranked factors associated with *E. coli* levels were site, dissolved oxygen, and avg. and min. air temperature (Fig. 1); site, dissolved oxygen, and avg. air temperature were among the four top-ranked factors regardless of the time period BSC considered when calculating the weather factors (Table S8). PDPs indicate that, on average, *E. coli* levels in the AZ canals (i) decreased as dissolved oxygen increased from 7 to 10 mg/L, and (ii) increased as avg. and min. air temperature 0-5 d BSC increased from 13°C to 33°C and from 3°C to 24°C, respectively (Fig. S7). For NY streams, the top-ranked factors associated with *E. coli* levels were turbidity, flow rate, pH, and min. air temperature; turbidity, flow rate, and pH were top-ranked factors regardless of the time period BSC considered (Table S8). On average, *E. coli* levels in the sampled streams increased as (i) turbidity increased from 0 to 50 NTUs, (ii) flow rate increased from 0.0 to 1.0 m/s, and (iii) min. air temperature 0-5 d BSC increased from 5°C to 18°C (Fig. S6). On average, *E. coli* levels in NY decreased as pH increased from 7.0 to 8.5 (Fig. S7).

### L. monocytogenes in AZ and NY

*L. monocytogenes* was isolated from 4% (3/76) of AZ GS and 15% (27/181) of NY GS (Table 1). While *L. monocytogenes* was isolated from 0 of the 34 AZ MS, *L. monocytogenes* was isolated from 7% (6/86; Table 2) of NY MS. In total, *L. monocytogenes* was isolated from samples collected on 29 of the 117 sampling days where paired MS-GS were collected (Table 3). *L. monocytogenes* was detected by MS only on 5 sampling days (all paired GS were *L. monocytogenes-*negative) and by one or more paired GS but not by the MS on 23 sampling days (Table 3). According to generalized linear mixed modelling, the odds of isolating *L. monocytogenes* from MS was significantly lower than the odds of isolating *L. monocytogenes* from a paired GS [Odds Ratio (OR=0.39); 95% Confidence Interval (CI) = 0.16, 0.97].

Random forest analysis could not be performed to identify factors associated with *L. monocytogenes* isolation in AZ due to the low *L. monocytogenes* prevalence in AZ. For random forests where *L. monocytogenes* isolation from NY GS or NY MS was the outcome, the random forest with the highest Kappa score was based on weather 0-4 days BSC (Kappa = 0.06; Accuracy = 0.76) and 0-1 day BSC (Kappa = 0.37; Accuracy = 0.90), respectively (Table S4). Random forest analysis identified flow rate as a top-ranked factor associated with *L. monocytogenes* isolation from GS and MS in NY (Fig. 1, S8). While the likelihood of *L. monocytogenes* isolation from GS decreased as flow increased from 0.0 to 1.0 m/s, the likelihood of *L. monocytogenes* isolation from MS increased as flow increased from 0.0 to 1.0 m/s (Fig. S9). The other top-ranked factors associated with *L. monocytogenes* isolation from GS were year-day, rainfall 3-4 d BSC, and avg. relative humidity 0-4 d BSC (Fig. 1). Flow rate and year-day were among the top-ranked factors regardless of the time period BSC considered (Table S8). The likelihood of *L. monocytogenes* isolation from NY GS decreased from May to July and increased from July to September (Fig. S8). Additionally, the likelihood of *L. monocytogenes* isolation from NY GS (i) increased as rainfall 3-4 d BSC increased from 0.0 to 2.0 cm, and (ii) decreased as avg. relative humidity 0-4 d BSC increased from 60% to 100% (Fig. S9). For NY MS the other top-ranked factors associated with *L. monocytogenes* isolation were pH, and min. and max. air temperature 0-1 d BSC (Fig. S8-S9).

**Figure 1:**
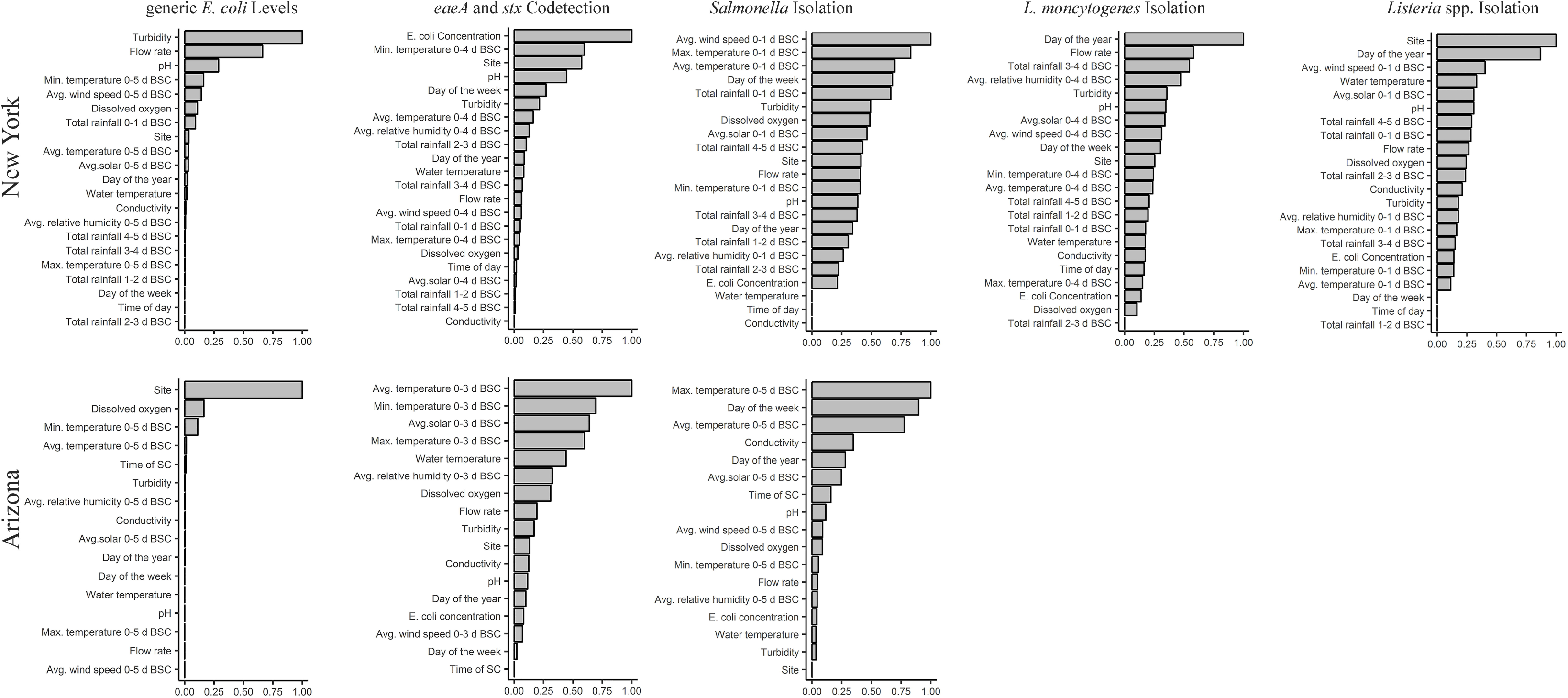
Results of random forest analyses that identified factors associated with *E. coli* levels, and the likelihood of codetecting *eaeA* and *stx*, and detecting *Salmonella, L. monocytogenes,* and *Listeria* spp. (including *L. monocytogenes*) in GS. The y-axis shows the factors ranked from most to least important. The x-axis shows NVI; a higher NVI (relative to all factors in the plot) equates to a stronger association between outcome and factor. NVI ≤ 0 indicates no association. Five, overlapping time frames (0-1, 0-2, 0-3, 0-4, or 0-5 d BSC) were used to calculate the values of the weather factors with the exception of rainfall; rainfall was calculated on a daily basis (0-1, 1-2, 2-3, 3-4, 4-5 d BSC). Separate forests were then developed for each outcome (e.g., *E. coli* levels in AZ, likelihood of *Salmonella* isolation in NY) and time frame in each state. The results for the forest with the highest Kappa score for each outcome are reported here. Thus, the time frame for the forest reported here differs for each combination of outcome and state. BSC = before sample collection.

**Table 4:** Results of generalized linear mixed models that characterized the relationship between the log_10_ 1383 MPN of *E. coli* level/100-mL and pathogen detection in grab samples.

| Target | Fixed Effects |  |  | Variance of Random Effects (SD<sup>c</sup>) |  |
| --- | --- | --- | --- | --- | --- |
|  | Change in Odds<sup>a</sup> | 95% CI<sup>b</sup> | *P*-value | Year-day | Site |
| *L. monocytogenes* |  |  |  |  |  |
| New York | 1.13 | 0.46, 2.77 | 0.786 | 0.5 (0.7) | 0.0 (0.0) |
| *Listeria* spp.<sup>d</sup> |  |  |  |  |  |
| New York | 0.97 | 0.37, 2.52 | 0.943 | 2.3 (1.5) | 1.3 (1.2) |
| *Salmonella* |  |  |  |  |  |
| Arizona | 4.43 | 1.46, 13.47 | 0.009 | 2.1 (1.4) | 0.0 (0.0) |
| New York | 1.53 | 0.71, 3.30 | 0.274 | 1.3 (1.1) | 0.2 (0.4) |
| *eaeA* and *stx* |  |  |  |  |  |
| Arizona | 50.20 | 4.05, 621.88 | 0.002 | 12.1 (3.5) | 6.6 (2.6) |
| New York | 18.40 | 5.39, 62.86 | < 0.001 | 1.1 (1.0) | 0.1 (0.3) |

<sup>a</sup> Change in the odds of detecting the target organism for a log<sub>10</sub> increase in the *E. coli* concentration.
<sup>b</sup> 95% Confidence Interval
<sup>c</sup> Standard Deviation
<sup>d</sup> Includes *L. monocytogenes*.

### *Listeria* spp. in AZ and NY

*Listeria* spp. (including *L. monocytogenes*) was isolated from 4% (3/76) of AZ GS and from 47% (85/181) of NY GS (Table 1). *Listeria* spp. was also isolated from 0 of the 34 AZ MS and 27% (23/86; Table 2) of NY MS. *Listeria* spp. was detected by MS only on 7 sampling days, and by one or more paired GS but not by the MS on 38 sampling days (Table 3). According to generalized linear mixed modelling, the odds of isolating *Listeria* spp. from MS was significantly lower than the odds of isolating *Listeria* spp. from a paired GS (OR=0.24; 95% CI=0.12, 0.48).

Random forest analysis could not be performed to identify factors associated with *Listeria* spp. isolation in AZ due to the low prevalence of *Listeria* spp. in AZ. For random forests where *Listeria* spp. isolation from NY GS or MS was the outcome, the random forest with the highest Kappa score was based on weather 0-1 days BSC (Kappa = 0.36; Accuracy = 0.69) and 0-5 days BSC (Kappa = 0.35; Accuracy = 0.75), respectively (Table S4). Random forest analysis identified site, year-day, avg. wind speed 0-1 d BSC and water temperature as the top-ranked factors associated with *Listeria* spp. isolation from NY GS (Fig. 1); site and year-day were among the 4 top-ranked factors regardless of the time frame BSC considered (Table S8). The likelihood of *Listeria* spp. isolation showed limited variation from May to July but increased from July to September (Fig. S9). Additionally, the likelihood of *Listeria* spp. isolation from GS (i) increased as the avg. wind speed 0-1 d BSC increased from 0 to 15 km/h, and (ii) decreased as water temperature increased from 10 to 23°C (Fig. S9). For NY MS the top-ranked factors associated with *Listeria* spp. isolation were rainfall 0-1 d BSC, min. and avg. air temperature 0-5 d BSC, and flow rate (Fig. S8).

### *Salmonella* in AZ and NY

*Salmonella* was isolated from 34% (26/77) of GS and 64% (21/33) of MS collected in AZ, and from 44% (80/181) of GS and 57% (50/88) of MS collected in NY (Tables 1-2. *Salmonella* was detected by the MS only on 30 sampling days, and by 1 or more paired GS but not by the MS on 21 sampling days (Table 3). The odds of isolating *Salmonella* from MS were 2.2 times greater than the odds of isolating *Salmonella* from a paired GS (OR=2.2; 95% CI=1.4, 3. 5).

For random forests where *Salmonella* isolation from AZ GS or MS was the outcome, the random forest with the highest Kappa score was based on weather 0-5 days BSC (Kappa = 0.40; Accuracy = 0.72) and 0-3 days BSC (Kappa = 0.38; Accuracy = 0.84), respectively (Table S5). According to random forest analysis, the two top-ranked factors associated with *Salmonella* isolation from AZ GS and MS were avg. and max. air temperature (Figs 1, S8). The likelihood of *Salmonella* isolation from AZ GS (i) increased as avg. and max. air temperature increased from 13 to 30°C and from 20 to 41°C, respectively, and (ii) decreased as avg. and max. air temperature increased from 30 to 38°C and from 41 to 47°C, respectively (Fig S10). The other top-ranked factors associated with *Salmonella* isolation from AZ GS were day of the week and conductivity; max. air temperature and day of the week were among the 4 top-ranked factors regardless of time period BSC considered (Table S8). The likelihood of isolating *Salmonella* from AZ GS was highest for samples collected on Tues. and Wed. (Fig. S10). Additionally, the likelihood of isolating *Salmonella* from AZ GS increased as conductivity increased from 750 to 1,300 uS/cm (Fig. S10).

**Table 5:**
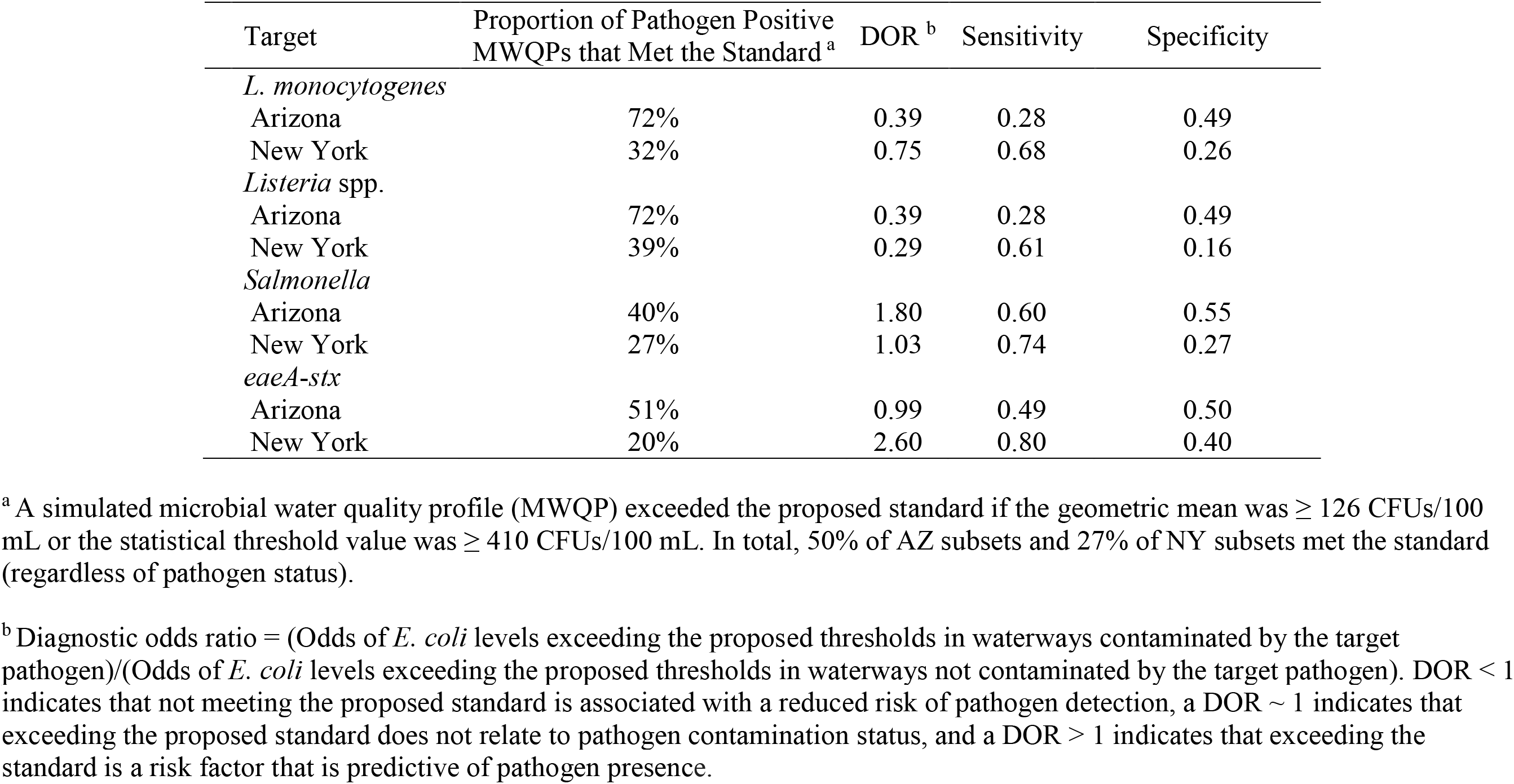
Ability of the proposed FSMA agricultural water standard (Food and Drug 1392 Administration, 2015) to predict *L. monocytogenes*, and *Listeria* spp., *Salmonella* isolation, and *eaeA-stx* codetection in agricultural water for a simulated dataset generated using a bootstrapping method.

For random forests where *Salmonella* isolation from NY GS or MS was the outcome, the random forest with the highest Kappa score for both GS BSC (Kappa = 0.18; Accuracy = 0.61) and MS BSC (Kappa = 0.11; Accuracy = 0.58) was based on weather 0-1 d BSC (Table S5). According to random forest analysis, the top-ranked factors associated with *Salmonella* isolation from NY GS were avg. wind speed 0-1 d BSC, max and avg. air temperature 0-1 d BSC, and day of the week (Fig. 1). The likelihood of *Salmonella* isolation from NY GS decreased as (i) avg. wind speed increased from 0 to 4 km/h, and (ii) avg. and max. air temperature increased from 10 to 19°C and from 15 to 26°C, respectively (Fig. S10). The likelihood of *Salmonella* isolation from NY GS increased as (i) avg. wind speed increased from 4 to 13 km/h, and (ii) avg. and max. air temperature increased from 19 to 26°C and from 26 to 33°C, respectively (Fig. S10). The likelihood of isolating *Salmonella* from NY GS was highest for samples collected on Sat. and lowest for samples collected on Wed. (Fig. S10). For NY MS the top-ranked factors associated with *Salmonella* isolation were rainfall 3-4 and 4-5 d BSC, turbidity, and year-day (Fig. S10).

### Codetection of *eaeA* and *stx* in AZ and NY

Forty-eight percent (44/83) of GS and 91% (30/33) of MS collected in AZ, and 69% (125/181) of GS and 88% (77/88) of MS collected in NY were PCR-screen positive for both *eaeA* and *stx* (Tables 1 & 2). Both genes were detected by MS only on 23 sampling days, and by 1 or more paired GS but not by MS on 7 sampling days (Table 3). The odds of codetecting *eaeA* and *stx* in a MS was 6.5 times greater than the odds of codetecting *eaeA* and *stx* in a paired GS (OR = 6.5; 95% CI= 3.1, 13.6).

While random forest analysis could not be performed to identify factors associated with *eaeA*-*stx* codetection in AZ MS due to the limited number of *eaeA* and *stx-*negative MS, random forest analysis was performed to identify factors associated with *eaeA*-*stx* codetection in AZ GS. For random forests where *eaeA*-*stx* codetection in AZ GS was the outcome, the random forest with the highest Kappa score was based on weather factors 0-3 days BSC (Kappa = 0.47; Accuracy = 0.74; Table S7). The top-ranked factors associated with *eaeA*-*stx* codetection in AZ GS were avg. solar radiation 0-3 d BSC, and avg., max., and min., air temperature (Fig. 1); avg. and max. air temperature were among the 4 top-ranked factors regardless of the time period BSC considered (Table S8). The likelihood of *eaeA*-*stx* codetection in AZ increased as (i) avg. solar radiation 0-3 d BSC increased from 11 to 31 Ly, and (ii) avg., max. and min. air temperature 0-3 d BSC increased from 10 to 27°C, from 20 to 42°C, and from 1 to 18°C, respectively (Fig. S11). The likelihood of *eaeA*-*stx* codetection in AZ decreased as avg. air temperature and min. air temperature 0-3 d BSC increased from 27 to 36°C and from 18 to 28°C, respectively (Fig. S11).

For random forests with *eaeA*-*stx* codetection in NY GS and MS as the outcome, the random forest with the highest Kappa score was based on weather factors 0-4 days BSC (Kappa = 0.52; Accuracy = 0.79) and 0-2 days BSC (Kappa = 0.24; Accuracy = 0.79), respectively (Table S7). The top-ranked factors associated with *eaeA*-*stx* codetection in NY GS were pH, min. air temperature 0-4 d BSC, the MPN of *E. coli*/100 mL, and site (Fig. 1); pH, *E. coli* levels, and site were among the 4 top-ranked factors regardless of the time period BSC considered (Table S8). The likelihood of *eaeA*-*stx* codetection in the NY GS increased as (i) min. air temperature 0-4 d BSC increased from 5 to 15°C, and (ii) *E. coli* levels increased from 18 to 1,000 MPN/100-mL (Fig. S11). The likelihood of *eaeA*-*stx* codetection in the NY GS decreased as (i) pH increased from 7.4 to 8.8, and (ii) min. air temperature 0-4 d BSC increased from 15 to 18°C (Fig. S11). The top-ranked factors associated with codetecting *eaeA* and *stx* in NY MS were rainfall 3-4 d BSC, conductivity, flow rate, and avg. air temperature 0-2 d BSC (Fig. S8).

### Effect of two-way interactions on microbial water quality

Due to the number of potential interactions that could have been investigated (e.g., 136 interactions could have been investigated for each random forest), we focused on the impact of biologically plausible interactions on estimated *E. coli* levels, and likelihood of detecting pathogens in GS (see Table S9 for a complete list). We focused on GS as opposed to MS because (i) approx. twice as many GS (N=264) were collected as MS (N=121), and (ii) GS are more commonly used to monitor surface water quality. Although the PDPs show evidence of threshold effects (e.g., stark differences in likelihood of detection above versus below a cut-off for a given factor), this may be a product of pseudoreplication, the sample size, the limited time span of the study, and/or the existence of true thresholds. Investigating these threshold effects is outside the scope of the current study. As such the results of the PDPs need to be interpreted with caution.

We found evidence of interactions between multiple factors (Figs S12-S19). For example, the likelihood of isolating *Salmonella* from AZ GS appeared to be higher when dissolved oxygen was <8.5 mg/L and air temperature was >20°C compared to when dissolved oxygen was >8.5 mg/L or air temperature was <20°C (Fig. S12). Similarly, estimated *E. coli* levels in AZ were higher when dissolved oxygen was <8.0 mg/L and air temperature was >28°C (Fig. S12-S13) compared to when dissolved oxygen was >8.0 mg/L or air temperature was <28°C. In AZ we also observed a synergistic interaction effect on likelihood of *Salmonella* isolation and likelihood of *eaeA*-*stx* codetection between dissolved oxygen and solar radiation, and between dissolved oxygen and water temperature (Fig S12).

We also found evidence of two-way interactions between turbidity and other factors (Fig. S14-S19). For instance, *E. coli* levels in NY were highest when rainfall 0-1 d BSC was >1 cm and turbidity was >10 NTU compared to when rainfall 0-1 d BSC was <1 cm or turbidity was <10 NTU (Fig S14). In NY we also observed a synergistic interaction effect between turbidity and (i) rainfall 0-1 d BSC on the likelihood of isolating *Salmonella*, and (ii) flow rate on estimated *E. coli* levels (Fig S15). Unlike the enteric targets, an antagonistic interaction effect on likelihood of *L. monocytogenes* isolation was observed for turbidity and rainfall 0-1 d BSC, and turbidity and flow rate (Fig S15).

Interactions between *E. coli* levels and other factors also appear to affect likelihood of pathogen detection (Fig. S16-S19). For instance, in AZ, likelihood of *Salmonella* isolation was lowest when *E. coli* levels were < 200 MPN/100-mL and avg. air temperature was < 20^◦^C, compared to when *E. coli* levels were > 200 MPN/100-mL or avg. air temperature was > 20^◦^C (Fig. S17). The likelihood of isolating *Salmonella* in NY GS appeared to be highest when *E. coli* levels were >1,350 MPN/100-mL and turbidity was >30 NTUs compared to when turbidity was <30 NTUs or *E. coli* levels were <1,350 MPN/100-mL (Fig. S18).

### Relationship between *E. coli* levels and pathogens in GS

The relationship between the log_10_ MPN of *E. coli*/100 mL and pathogen detection was characterized using generalized linear mixed models. Models could not be developed to characterize the relationship between *Listeria* isolation and *E. coli* levels in AZ due to the low prevalence of *Listeria* in AZ. According to these analyses, *Salmonella* isolation in AZ, and *eaeA-stx* codetection in AZ and NY were significantly associated with *E. coli* levels, but *L. monocytogenes* and *Salmonella* isolation in NY were not (Table 4). The odds of isolating *Salmonella* from AZ GS increased by a factor of 4 (95% CI = 1.5, 13.5) for each log_10_ increase in the MPN of *E. coli*/100-mL. The odds of codetecting *eaeA* and *stx* in AZ and NY GS increased by a factor of 50 (95% CI = 4.1, 621.9) and a factor of 18 (95% CI = 5.4, 62.9), respectively, for each log_10_ increase in the MPN of *E. coli*/100-mL.

We also assessed the predictive accuracy of the proposed FSMA standard [geometric mean <126 CFUs/100-mL and STV <410 CFUs/100-mL; (Food and Drug Administration, 2015)] to identify waterways with a high or low risk of pathogen presence at time of water use. Briefly, bootstrapping was used to simulate water sampling to create a microbial water quality profile (MWQP) composed of 20 samples (N=10,000 MWQPs per waterway). The last GS selected for inclusion in each MWQP represented water quality at time of water use. The geometric mean and STV varied substantially among the simulated MWQPs for a given waterway (Fig. 2). While approx. 50% of MWQPs in AZ and 27% of MWQPs in NY met the proposed FSMA standard (Fig. 2; Table 5), the percent of pathogen-positive MWQPs that met the standard ranged between 20% (*eaeA-stx* codetection in NY) and 72% (*L. monocytogenes* in AZ). In general, the efficacy of the proposed standard for identifying waterways contaminated by pathogens appears to be region and pathogen-specific. For instance, while the odds of *E. coli* levels exceeding the standard was 2.6 times greater for streams positive *eaeA* and *stx* compared to streams negative for both genes [diagnostic odds ratio (DOR) = 2.6], the odds of *E. coli* levels exceeding the standard were approx. equal for canals positive *eaeA* and *stx*, and for canals negative for both genes (DOR=0.99). We also found that the odds of *E. coli* levels exceeding the standard was lower for *L. monocytogenes*-positive waterways compared to *L. monocytogenes*-negative waterways (DOR=0.4 in AZ; DOR=0.8 in NY).

**Figure 2:**
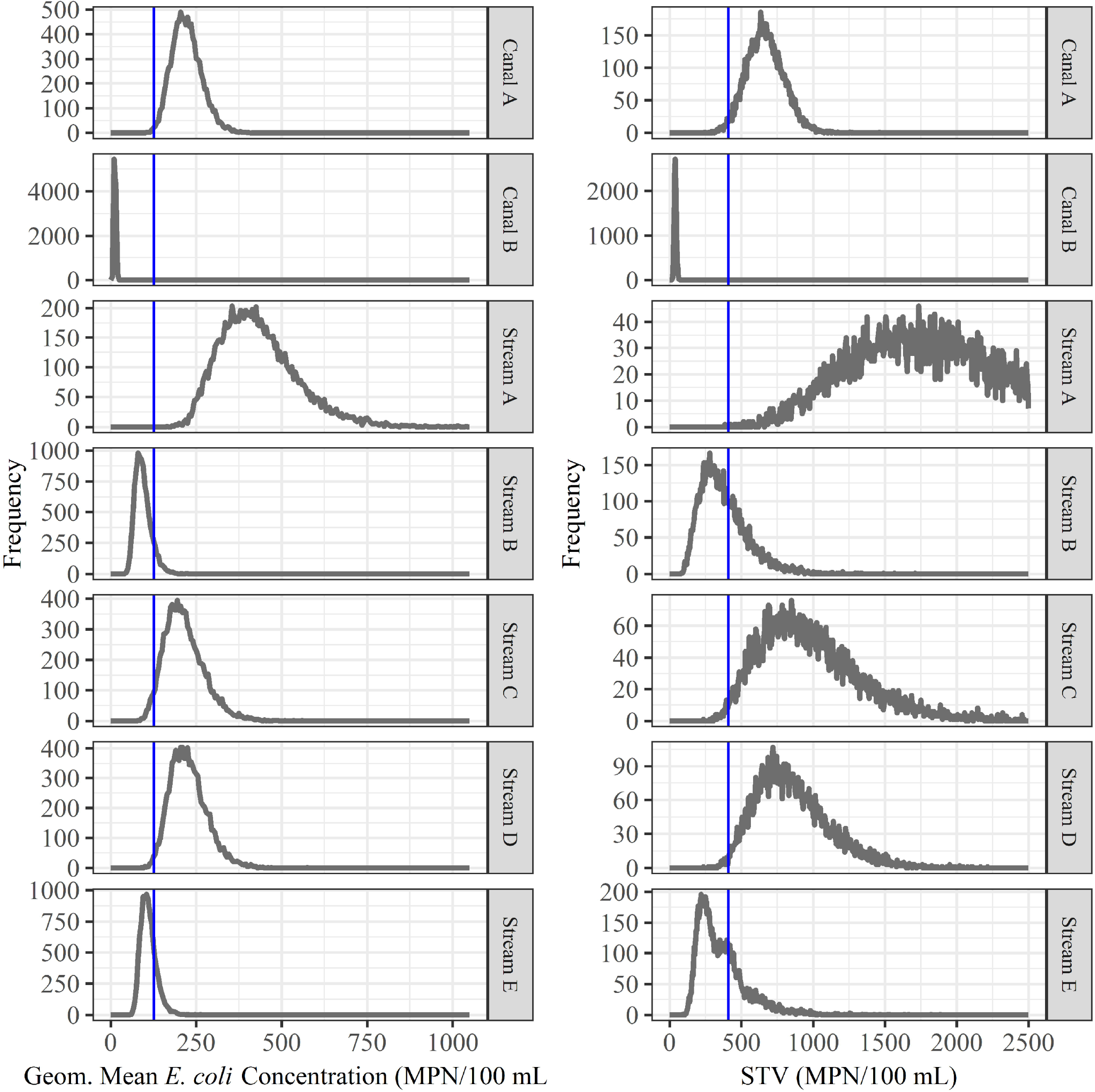
Bootstrapping was used to simulate water sampling to create microbial water quality profiles (MWQP) composed of 20 samples (N=10,000 MWQPs per waterway). The graphs show the geometric mean and statistical threshold value (STV) for all MWQPs for each waterway. The blue line represents the proposed FSMA standard cut-offs [geometric mean < 126 CFU/100 mL and STV < 410 CFU/100 mL; (Food and Drug Administration, 2015)]. Note that the y-axis varies between plots.

## DISCUSSION

Management of agricultural water-associated microbial food safety risks is not amenable to one-size-fits-all approaches due to the heterogeneous nature of freshwater environments and as environmental conditions affect both the risk of pathogen detection and the relationship between indicator organism levels and foodborne pathogen presence. To improve our ability to develop location-specific management practices, this study was conducted in two US produce-growing regions to (i) characterize the relationship between *E. coli* levels and pathogen presence in agricultural water, and (ii) identify environmental factors associated with pathogen detection. This study is unique in its use of advanced machine learning approaches (e.g., random forest, partial dependence plots) to examine the impact of multiple interactions between environmental factors on the likelihood of detecting foodborne pathogens or pathogen markers (i.e., the *eaeA* and *stx* genes) in agricultural water. In addition, this study provided data on several understudied topics, including (i) produce safety risks associated with preharvest surface water use in New York state, and (ii) the recovery of *Listeria* from surface water sources using More swabs (MS) compared to 10-L grab samples (GS) filtered through modified Moore swabs (mMS).

Overall, our study not only showed that sampling methods can affect reported pathogen prevalence, but also indicated that interactions between environmental factors (e.g., rainfall, turbidity) (i) were associated with likelihood of pathogen detection, and (ii) mediated the relationship between *E. coli* levels and likelihood of pathogen detection. Our findings suggest that (i) environmental heterogeneity, including interactions between environmental factors, affect microbial water quality, and (ii) *E. coli* levels alone may not be a suitable indicator of the food safety risks associated with preharvest water use. Alternative methods that utilize environmental and microbial data (e.g., monitoring programs that use physiochemical and microbial water quality data to identify when and where foodborne pathogens may be present in agricultural waterways) are needed to assess the food safety risks associated with preharvest water use. However, the findings reported here have to be viewed in the context of several limitations. For instance, the data reported here were collected from nine waterways over one growing season. Additionally, while the NY streams were randomly selected from the total population of eligible streams, the AZ canals were enrolled based on convenience for the sampling team, which means that the results reported here may be affected by selection bias. For instance, a cattle feedlot was immediately upstream and next to the sampling site for Canal A, while feedlots were not present near Canals B and C. This could explain why Canal A had significantly higher *E. coli* levels compared to Canals B and C. *Even though the pathogen prevalence and E. coli levels reported here may not be generalizable to other AZ canals, our study highlights the variability in microbial water quality in AZ, and the resulting limitations associated with the use of a single microbial indicator to guide management of the food safety risks associated with preharvest surface water use.* These data, when taken in aggregate with data from other ongoing or recent studies, will facilitate the development of science-based strategies for managing the food safety risks associated with preharvest surface water use. Additionally, since this is an observational study, the data reported here cannot be used to establish causal relationships but instead indicate potential associations. Since water contamination is frequently a stochastic process, the identification of associations here may be due to random chance as opposed to true associations. Thus, additional studies are needed to determine if our findings are generalizable for the sampled waterways in future years, and for other waterways in AZ, NY, and other regions. While a strength of this study is its longitudinal nature, this also resulted in pseudo-replication and potential autocorrelation. To address these concerns, site and year-day were included as random effects or covariates in all analyses. Despite the aforementioned limitations, our findings indicate that approaches that account for temporal variation in environmental conditions, including microbial water quality, are needed to help identify and address food safety risks associated with the preharvest use of surface water for produce production.

### Reported pathogen prevalence differed when different sampling methods were used

One objective of this study was to compare the ability of 24-h MS and 10-L GS to detect foodborne pathogens in surface water. Conceptually, a GS acts as a snapshot and provides data on water quality at a specific time, while MS capture bacteria that flow through the waterway over a given time period (24 h in our study). We therefore hypothesized that MS would be better than GS at detecting pathogens. As predicted, the likelihood of *Salmonella* isolation, and *eaeA-stx* codetection was significantly greater for MS compared to GS. Past studies that compared the ability of MS and GS to detect pathogens in surface water reported similar results (Benjamin et al., 2013; Colburn et al., 1990; Hoganson and Elliott, 1972). A study that used MS and 100-mL GS to sample California waterways found that the proportion of *E. coli* O157:H7-positive MS (13.8%; 12/87) was significantly greater than the proportion of O157:H7-positive GS [1.8%; 10/558; (Benjamin et al., 2013)]. Unlike *Salmonella* isolation and *eaeA*-*stx* detection, the likelihood of *Listeria* isolation was significantly lower for MS compared to GS in our study. A potential explanation for the lower than expected rate of *Listeria* detection by MS is that competitive microflora in the MS inhibit *Listeria* recovery. In fact, we observed more competitive microflora and fewer *Listeria*-like colonies when plating MS enrichments compared to paired GS enrichments. Past studies (Buzoleva and Terekhova, 2002; Francis and O’Beirne; Habimana et al., 2011; Hansen et al., 2006; Locatelli et al., 2013; McLaughlin et al., 2011), which found that the presence of other microflora inhibited *Listeria* survival, support this hypothesis. For instance, McLaughlin et al. (McLaughlin et al., 2011) studied *L. monocytogenes* survival in soil and found that *L. monocytogenes* populations grew in the absence of other microflora but rapidly declined when other microflora were present. *Salmonella* recovery may not have been affected by competitive microflora because samples used for *Salmonella* isolation underwent multiple selective, enrichment steps while samples used for *Listeria* isolation underwent one selective, enrichment step. These findings indicate that appropriate water sampling methods depend on the reason for sample collection. Specifically, the organism of concern (e.g., AZ leafy green growers concerned about pathogenic *E. coli* may decide to use MS as opposed to GS to test water sources), study aims (e.g., is the goal to optimize pathogen detection or replicate industry practices), time constraints (MS require 2 site visits but GS require 1 visit), outcome of interest (e.g., MS cannot be used to calculate concentrations since the volume of water that flows through a MS is unknown), and potential loss of MS to storms or human tampering should be considered when selecting MS or GS as a sampling method.

In examining pathogen prevalence in the current study, it is important to note that the prevalence of *L. monocytogenes* in the current study was substantially lower than the prevalence reported by past NY studies; the current study used the same *Listeria* enrichment and isolation protocols as these past studies (Chapin et al., 2014; Strawn et al., 2013a, 2013b; Weller et al., 2015b, 2015a). For instance, we isolated *L. monocytogenes* from 6% (2/32) of 10-L GS collected from Stream C, while two studies that sampled Stream C at approx. the same site as our study isolated *L. monocytogenes* from 71% (15/21; unpublished) and 63% [33/52; (Weller et al., 2015a)] of 250-mL GS in 2013 and 2014, respectively. Collection of a larger volume of water in the current study required a change in the GS filtration protocol; instead of filtering 250-mL GS through a 0.45 um filter like previous studies (Chapin et al., 2014; Strawn et al., 2013a, 2013b; Weller et al., 2015b, 2015a), we filtered 10-L through a mMS. We speculate that the lower than expected *Listeria* prevalence in the current study is because the mMS-method has not been optimized for *Listeria* recovery. Indeed, while several studies examined *Salmonella* and *E. coli* recovery from large volumes of water using mMS (Bisha et al., 2011, 2014; McEgan et al., 2013b; Sbodio et al., 2013; Zhu et al., 2019), no study, to our knowledge, has quantified *Listeria* recovery using mMS. Such a study is needed if the mMS approach is to be incorporated into industry and government water testing programs as previously suggested (Bisha et al., 2014).

### Microbial water quality varied across time and space

Random forest analysis identified associations between temporal factors and (i) *E. coli* levels, and (ii) the likelihood of pathogen detection. For example, year-day (i.e., number of days since Jan. 1, 2017) was associated with detection of all target organisms from both sample types in both states and was among the top-ranked factors associated with *L. monocytogenes*, and *Listeria* spp. isolation in NY GS. The identification of an association between year-day and microbial water quality suggests that microbial water quality varied seasonally, which is consistent with the findings of previous studies (Carter et al., 1987; Cooley et al., 2014; Falardeau et al., 2017; Gorski et al., 2011; Horman et al., 2004; Wilkes et al., 2009). For example, Falardeau et al. (Falardeau et al., 2017) reported that *L. monocytogenes* was more prevalent in winter than summer in agricultural watersheds in British Colombia, Canada. Similarly, Cooley et al. (Cooley et al., 2014) found that the likelihood of isolating *L. monocytogenes* from California surface water samples was significantly higher in winter and spring compared to summer and fall. Falardeau et al. (Falardeau et al., 2017) and Cooley et al. (Cooley et al., 2014) attributed these seasonal trends in *L. monocytogenes* isolation to increased rainfall and lower temperatures in winter and spring compared to summer and fall. In our study, the likelihood of isolating *L. monocytogenes* in NY (i) was also associated with temperature, and (ii) decreased as air temperature increased from May to June and increased as air temperature decreased from July to September. However, year-day may also serve as a proxy for unmeasured factors that change seasonally and affect microbial water quality, including anthropogenic reductions in water flow (e.g., damming of irrigation ditches), which inhibit pathogen movement within watersheds (Falardeau et al., 2017). While we did not observe this type of activity, all of the sampled waterways are in agricultural areas and provide water to commercial operations. As such, upstream activity that varied over the course of the growing season may have contributed to the seasonal variation in microbial water quality discussed above.

Microbial water quality and likelihood of pathogen detection also varied between waterways in our study. Sample site, which is unique to each waterway sampled here, was among the four top-ranked factors associated with *E. coli* levels in AZ, and with detecting *Listeria* spp. and codetecting *eaeA* and *stx* in NY GS. Given the number of factors (e.g., bottom substrate, upstream land use) that differ between waterways and may affect water quality, it is not surprising that microbial water quality would vary between waterways. In fact, multiple studies have found associations between microbial water quality and upstream land use (Bradshaw et al., 2016; Brendel and Soupir, 2017; Dila et al., 2018; Falardeau et al., 2017; Lyautey et al., 2007; Pandey et al., 2012; Verhougstraete et al., 2015). For instance, Pandey et al. (Pandey et al., 2012) tracked water quality at 46 sites in an Iowa watershed and found that *E. coli* levels were positively associated with the amount of cropland in Thiessen polygons around each site. Associations between microbial water quality, and proximity to upstream livestock operations (Bond and Partyka, 2004; Lyautey et al., 2010; Wilkes et al., 2011), the number of septic systems in a watershed (Verhougstraete et al., 2015), and livestock and human population density (Dila et al., 2018; Falardeau et al., 2017) have also been reported. Post-hoc identification of factors that drive spatial variation in water quality is difficult and requires data that were not collected as part of the current study (e.g., on upstream activity and land use). Despite this limitation, our findings indicate that microbial water quality varies between waterways. Future studies should therefore consider the impact of spatial factors, in addition to weather and physiochemical water quality, when investigating associations between environmental conditions and microbial water quality.

### Weather and physiochemical water quality were associated with *E. coli* levels as well as the likelihood of detecting foodborne pathogens in NY and AZ surface water

Although the 4 top-ranked factors associated with *E. coli* levels in AZ were site, DO, and avg. and min. air temperature, the variable importance scores (VI) for site and dissolved oxygen were substantially larger than the VI scores for avg. and min. air temperature. Similarly, the VI scores for turbidity, flow rate and pH, the 3 top-ranked factors associated with *E. coli* levels in NY, were substantially greater than the score for min. air temperature, the 4^th^-ranked factor. This suggests that *E. coli* levels were more strongly associated with physiochemical water quality than weather for the waterways sampled during our study. Previous studies have repeatedly identified associations between physiochemical water quality and *E. coli* levels (Ansa et al., 2011; Christensen et al., 2000; Horman et al., 2004; Rao et al., 2015; Roslev et al., 2004; Stocker et al., 2016). Similar to our study, a study that examined the microbial and physiochemical quality of three Ecuadorian rivers found a negative association between dissolved oxygen and *E. coli* levels (Rao et al., 2015). A potential explanation for the inverse relationship between dissolved oxygen and *E. coli* levels is that the ability of UV radiation to damage bacterial cells is positively associated with dissolved oxygen (Ansa et al., 2011; Curtis et al., 1992; Davies-Colley et al., 1997; Ouali et al., 2014). While this hypothesis is not supported by the interaction between solar radiation and dissolved oxygen observed in our study in AZ (estimated *E. coli* levels in AZ were highest when dissolved oxygen was low and solar radiation was high), it is supported by the interaction observed in NY (estimated *E. coli* levels in NY were highest when both dissolved oxygen and solar radiation were low). This discrepancy may be due to the correlation between solar radiation and temperature in AZ, which confounds the true nature of the interaction between dissolved oxygen and solar radiation.

While some studies have identified an association between rainfall and microbial water quality (Francy et al., 2013; Pandey et al., 2012; Stocker et al., 2016; Wilkes et al., 2011; Won et al., 2013), other studies did not (Benjamin et al., 2013; McEgan et al., 2013a; Won et al., 2013). For instance, Won et al. (Won et al., 2013) reported that *E. coli* levels in Ohio reservoirs were not associated with rainfall 0-1 d BSC, while *E. coli* levels in Ohio canals were higher on days when >2 cm of rain fell. Interestingly, a survey of Florida water sources, which did not find an association between *Salmonella* levels and rainfall over the day, week, or month preceding sampling, hypothesized that rainfall did not have a direct effect on microbial water quality, and instead interacted with other factors to affect microbial water quality (McEgan et al., 2013a). Our finding that synergistic interactions between rainfall and turbidity were associated with *E*. *coli* levels as well as the likelihood of detecting *Salmonella* and codetecting *eaeA* and *stx* in NY GS support this hypothesis. Rainfall and turbidity are both indicative of conditions that facilitate pathogen movement from environmental sources into the water column, which may explain the synergistic interaction observed here. For example, during rain events, run-off and flooding provide a pathway for pathogen movement from terrestrial sources into waterways. Similarly, stream sediments can act as an in-channel store of bacteria, and disturbance of these sediments during rain events can re-introduce bacteria into the water column and concomitantly elevate turbidity levels (Jamieson et al.; Muirhead et al., 2004; Nagels et al., 2002). However, due to the correlation between environmental factors in our study, determining the exact nature of the interactions observed (e.g., if temperature, solar radiation or both factors in concert interact with dissolved oxygen to affect *E. coli* levels in AZ) requires additional data not collected here and is therefore beyond the scope of the present study. Despite this limitation, our findings suggest temporal environmental heterogeneity (i.e., interactions between environmental factors and changes in environmental conditions over time) is associated with microbial water quality and should be taken into account when designing strategies for mitigating food safety risks associated with preharvest surface water use.

Even though our study found that complex interactions between weather and water quality factors were associated with microbial water quality, our findings also suggest relationships between specific factors and microbial water quality are reproducible. These factors may therefore be useful as supplemental indicators of microbial water quality. For example, turbidity was ranked as an important factor in 7 of the 15 random forest analyses performed here, and was associated detecting pathogens in samples collected from both AZ and NY waterways. Multiple studies (Christensen et al., 2000; Francy et al., 2013; Havelaar et al., 2017; Horman et al., 2004; Nagels et al., 2002; Rao et al., 2015; Topalcengiz et al., 2017), including the study reported here and the Ecuadorian study discussed above (Rao et al., 2015), found a positive association between *E. coli* levels and temperature, or between *E. coli* levels and turbidity. Francy et al. (Francy et al., 2013) surveyed recreational water quality at 22 Ohio beaches along inland lakes and found that turbidity was one of the best predictors of *E. coli* levels. Like the study presented here, previous studies have also found associations between turbidity and pathogen presence in surface water (Francy et al., 2013; Partyka et al., 2018; Wilkes et al., 2011). For instance, Wilkes et al. (Wilkes et al., 2011) and Partyka et al. (Partyka et al., 2018) developed models to predict the presence of foodborne pathogens in Ontario, and California and Washington surface water, respectively. Wilkes et al. (Wilkes et al., 2011) found that turbidity was an informative predictor of *E. coli* O157:H7 and *L. monocytogenes* presence, and that 93% of *E. coli* O157:H7 positive samples were collected when turbidity was > 4.8 NTUs (Wilkes et al., 2011). Partyka et al. (Partyka et al., 2018) found that the odds of isolating *Salmonella* increased 3-fold for each log_10_ increase in turbidity.

Although we identified a relationship between air and water temperature and microbial water quality in 14 of the 15 random forest analyses performed here, this relationship is complex and, as such, temperature may not be suitable as a supplemental indicator of microbial water quality. For instance, temperature was correlated with year-day and solar radiation among other factors, which obfuscates our ability to interpret the relationship between temperature and likelihood of pathogen detection. Moreover, based on the findings of this and other studies, the strength and direction of the relationship between temperature and microbial water quality appear to be pathogen, region, and/or waterway-specific (Francy et al., 2013; Luo et al., 2015; Truchado et al., 2018). In the study reported here, the likelihood of detecting *Salmonella* in AZ GS, and of codetecting *eaeA* and *stx* in NY MS increased as avg. air temperature increased from 15 to 20°C while the likelihood of *Salmonella* isolation from NY GS decreased over the same temperature range. Similarly, Francy et al. (Francy et al., 2013) found that the strength and direction of correlation between water temperature and *E. coli* levels in 22 Ohio lakes were different for different water sources in the same region. Overall, the findings from this and other studies suggest that turbidity but not temperature may be useful as a supplemental indicator of the microbial quality of surface water (Havelaar et al., 2017; Stocker et al., 2016). Our data do however suggest that temperature is strongly associated with microbial water quality, and inclusion of temperature as a factor in more sophisticated algorithms (e.g., AI tools, machine learning-based predictive risk maps) for predicting when and where water is likely to be contaminated by foodborne pathogens may be useful.

### The relationship between *E. coli* levels and pathogen detection in surface water appears to be mediated by environmental conditions

In the current study, we found that the relationship between *E. coli* levels and pathogen detection was region- and pathogen-specific. For example, we found that *E. coli* levels were significantly associated with *Salmonella* isolation in AZ but not in NY. These findings are not unexpected since the relationship between *E. coli* levels and pathogen presence in surface water varied widely between past studies (Benjamin et al., 2013; Economou et al., 2013; Harwood et al., 2005; McEgan et al., 2013a; Pachepsky et al., 2015; Wilkes et al., 2009). Pachepsky et al. (Pachepsky et al., 2015) compiled the findings of 40 datasets with information on *E. coli* levels and pathogen presence in surface water, and found evidence of a significant relationship between *E. coli* levels and pathogen detection in 18% of the datasets. Similarly, McEgan et al. (McEgan et al., 2013a) found that the relationship between *E. coli* and *Salmonella* levels varied substantially between 18 Florida waterways, and hypothesized that environmental factors mediated the relationship between *E. coli* and *Salmonella* levels in their study (McEgan et al., 2013a). Their hypothesis is supported by the findings of Bradshaw et al. (Bradshaw et al., 2016), who used classification trees to predict when Georgia waterways were contaminated by enteric pathogen,s and found that *E. coli* was a useful predictor only when certain conditions were met. Specifically, *E. coli* levels were useful for identifying (i) *Salmonella*-positive samples when dissolved oxygen <11.3 mg/L and pH <6.65, and (ii) *stx*-positive samples when air temperature ≥13°C (Bradshaw et al., 2016). These findings suggest that the relationship between *E. coli* levels and likelihood of pathogen presence may be weather, region, and/or pathogen-dependent. As such, the use of *E. coli* alone may not be a suitable indicator of different food safety risks associated with preharvest surface water use; this conclusion is consistent with those of other recent studies [e.g., (Havelaar et al., 2017; Truitt et al., 2018)].

Due to the continued use of *E. coli* to assess food safety risks associated with preharvest surface water use (California Leafy Greens Marketing Agreement, 2017; Food and Drug Administration, 2015; Jones and Shortt, 2010; Rock et al., 2018), understanding the associations between specific environmental factors and the relationship between *E. coli* levels and pathogen contamination of surface water is critical. As such, we used two-way PDPs to identify potential associations between the likelihood of detecting foodborne pathogens in AZ and NY waterways, and two-way interactions between environmental factors and *E. coli* levels. We found evidence that interactions between *E. coli* levels and multiple environmental factors, including dissolved oxygen and turbidity, were associated with the likelihood of detecting one or more pathogens. One of the basic tenets of ecology is that different organisms will respond differently to the same environmental conditions. One would therefore expect pathogens to respond to environmental conditions in a different manner than *E. coli* (e.g., *Salmonella* populations may persist while *E. coli* populations may die-off under a given condition). Indeed, multiple studies have shown that *E. coli* and foodborne pathogens respond differently to solar radiation (Jenkins et al., 2011; McCambridge and McMeekin, 1981; Sinton et al., 2007), temperature (Martinez et al., 2014; Rhodes and Kator, 1988), and other environmental conditions (Hood and Ness, 1982; Mezrioui et al., 1995). In fact, a review that compiled findings on *E. coli* and *Salmonella* survival in non-host environments concluded that *Salmonella* was able to survive under a wider variety of environmental conditions and persist for longer in aquatic environments than *E. coli* (Winfield and Groisman, 2003). Moreover, the variation in the relationship between *E. coli* levels and pathogen presence observed here and in other studies [e.g., (McEgan et al., 2013a)], may also be a product of the fact that sources of generic *E. coli* and specific pathogens may differ. For example, recent studies have found evidence that *E. coli,* including pathogenic *E.coli*, and *Salmonella* can exist as autochthonous or naturalized populations in non-host environments [e.g., water (Goto and Yan, 2011; Hendricks, 1967; McEgan et al., 2013a), algal mats (Byappanahalli et al., 2003; Ksoll et al., 2007; Whitman et al., 2003), soil (Goto and Yan, 2011; Ishii et al., 2010; NandaKafle et al., 2018; Nautiyal et al., 2010)]. Thus, the co-occurrence of *E. coli* and foodborne pathogens in surface water environments is not necessarily evidence of a recent fecal contamination event but may instead result from conditions that facilitate concomitant contamination from different sources, or the growth and survival of both organisms (McEgan et al., 2013a). Thus, our conclusion that environmental conditions mediate the relationship between *E. coli* levels and the likelihood of pathogen contamination of surface water sources is logical when viewed through the lens of ecology and in the context of the existing literature. By mediating the relationship between *E. coli* levels and pathogen presence in surface water, environmental conditions complicate interpretation of *E. coli*-based water test results. This further illustrates the challenges associated with identifying and using a single parameter like *E. coli* levels as the primary basis for making decisions on how to best mitigate food safety risks associated with preharvest surface water use for produce production.

### The proposed FSMA standard is not indicative of the food safety risks associated with preharvest surface water use

It is important to consider how water testing results are interpreted when examining the use of *E. coli* as an indicator of the food safety risks associated with preharvest surface water use. For example, the proposed FSMA standard states that growers must collect 20 samples over a 2 to 4 year period to create a microbial water quality profile [MWQP; (Food and Drug Administration, 2015)]. The geometric mean *E. coli* level and STV of the MWQP must be ≤126 CFUs/100 mL and ≤410 CFUs/100 mL, respectively (Food and Drug Administration, 2015). We found that the geometric mean and STV varied substantially among the simulated MWQPs for each waterway; for instance, the geometric mean of the simulated MWQPs for Stream E varied between 56 and 265 MPN/100-mL. This indicates that meeting the proposed FSMA standard is largely a function of when the water samples that comprise the MWQP were collected, and that meeting the standard may be a poor approximation of *E. coli* levels in surface water at the time of water use. Additionally, when we quantified the ability of the proposed standard to identify the pathogen status of the simulated MWQPs, we found that the predictive accuracy of the proposed standard was poor with regard to predicting (i) *eaeA-stx* codetection, or *L. monocytogenes* detection in AZ canal water, and (ii) *Salmonella* or *L. monocytogenes* detection in NY stream water (the DOR was less than or approx. 1). One limitation of the simulated sampling reported here is that our samples were collected over one growing season while the proposed standard uses samples collected over 2 to 4 years to create the MWQP. However, our conclusions are logical given the temporal variation in the microbial quality of surface water observed in this and other studies (Goyal et al., 1977; Hipsey et al., 2008; Pandey et al., 2012; Payment and Locas) as well as our finding that the relationship between *E. coli* levels and pathogen presence was mediated by environmental conditions. Moreover, our finding is consistent with that of Havelaar et al. (Havelaar et al., 2017) who also examined the predictive accuracy of the proposed FSMA water quality standard and found that MWQPs consisting of 20 samples were insufficient to capture the variability in *E. coli* concentrations in Floridian agricultural water sources. Overall these findings suggest that *E. coli* alone may not be a reliable indicator of the food safety risks associated with preharvest surface water use. The fact that other studies (Benjamin et al., 2013; Harwood et al., 2005; Havelaar et al., 2017; Ishii and Sadowsky, 2008; McEgan et al., 2013a; Pachepsky et al., 2015; Topalcengiz et al., 2017; Truitt et al., 2018) conducted out in other regions, in other years, and using different protocols reached the same conclusion as the study reported here, suggests that our conclusion is robust despite limitations associated with our study’s observational nature and time frame.

## Conclusion

Using advanced machine learning approaches this study showed that microbial water quality is associated with temporal environmental heterogeneity (interactions between environmental factors and changes in environmental conditions over time). As such, the food safety risks associated with preharvest use of a given surface water source are not constant over time and instead depend on environmental conditions at the time of water use. Our findings also indicate that (i) the relationship between *E. coli* levels and pathogen presence in surface water is mediated by environmental conditions, and (ii) *E. coli* levels alone may not be a suitable indicator of the food safety risks associated with preharvest surface water use. Instead, alternative approaches [e.g., e.g., models that incorporate data on *E. coli* levels and environmental conditions, incorporation of turbidity as a supplementary indicator into E. coli-based water quality monitoring programs] are needed to improve growers’ ability to identify and address these food safety risks in real-time. Given the dynamic and complex nature of surface water systems these alternative approaches need to (i) account for temporal variation in weather, and in physiochemical and water quality, and (ii) provide insights on microbial water quality at the time of water use.

## Author Contributions

DW and MW conceived of the project idea. DW, MW, and CR designed the study, and wrote the grant. DW, MW, CR and NB coordinated efforts between the NY and AZ teams. DW and NB oversaw day-to-day aspects of the project. DW, NB, SR, and EG carried out field and laboratory work. DW, EM, and RI developed the data analysis plan, which DW implemented. DW wrote the paper with input from all authors.

## Acknowledgments

This research was largely funded by a grant from the Center for Produce Safety under award number. Manuscript preparation was supported by the National Institute of Environmental Health Sciences of the National Institutes of Health (NIH) under award number T32ES007271. The content is solely the responsibility of the authors and does not represent the official views of the NIH.

We are grateful for the technical assistance of Maureen Gunderson, Alexandra Belias, and Deniz Akdemir. We are also grateful to Aziza Taylor, Kyle Markwardt, Sriya Sunil, Ahmed Gaballa, and Xiaodong Guo for help in the field and the laboratory.

## Abbreviations

BSC: before sample collection
CFU: colony-forming units
DOR: Diagnostic odds ratio
FSMA: Food Safety Modernization Act
GS: grab sample(s)
mMS: modified Moore swab(s)
MPN: most probable number
MS: Moore swab(s)
MWQP: microbial water quality profile(s)
NVI: normalized variable importance
PDPs: partial dependence plot(s)
STV: statistical threshold value
VI: variable importance

