## Supplemental Materials for "Complex interactions between weather, and microbial and physiochemical water quality impact the likelihood of detecting foodborne pathogens in agricultural water"

Figure S1: Samples were collected using a “1-week sampling” scheme and a “1-day sampling” scheme. For 1-week sampling (Fig A), samples were collected each day for up to 6 consecutive days. On the first day of 1-week sampling, a Moore swab (MS) set was anchored in the waterway, and a grab sample (GS) set were collected. When those MS were collected 24-h later, a second GS set was collected and three more MS were placed in the waterway. This was repeated daily for up to 6 d. During 1-day sampling, one MS was left in the waterway for 24-h, and 6 GS sets were collected between 6 am and 8 pm approx. 2.5 hours apart.

A

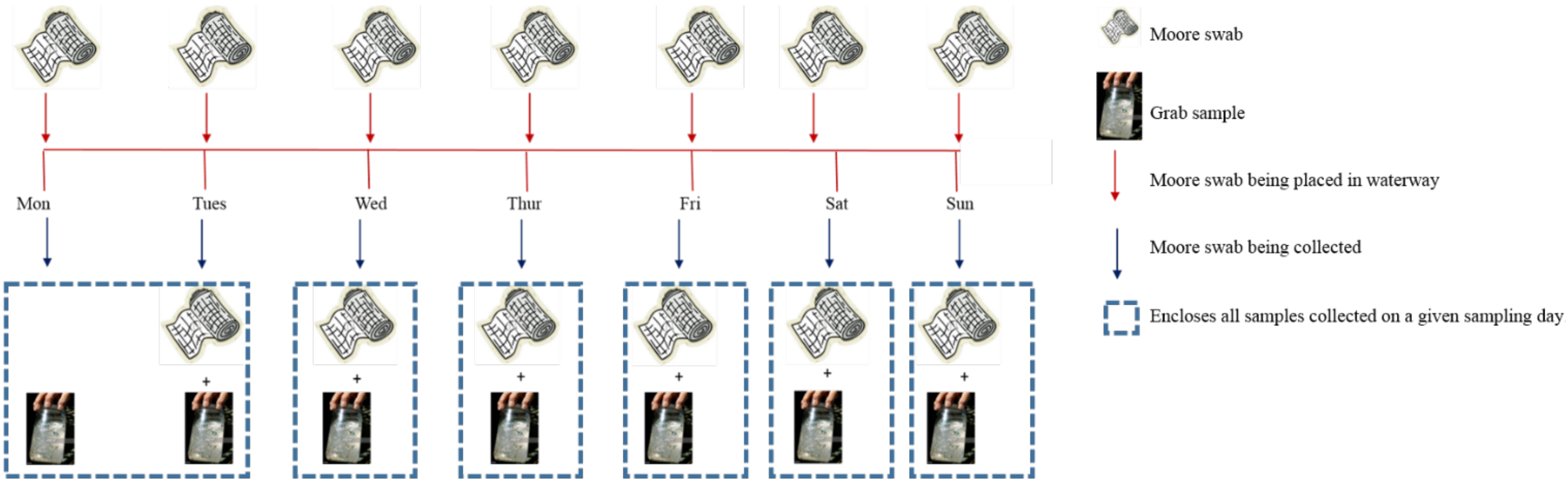


B

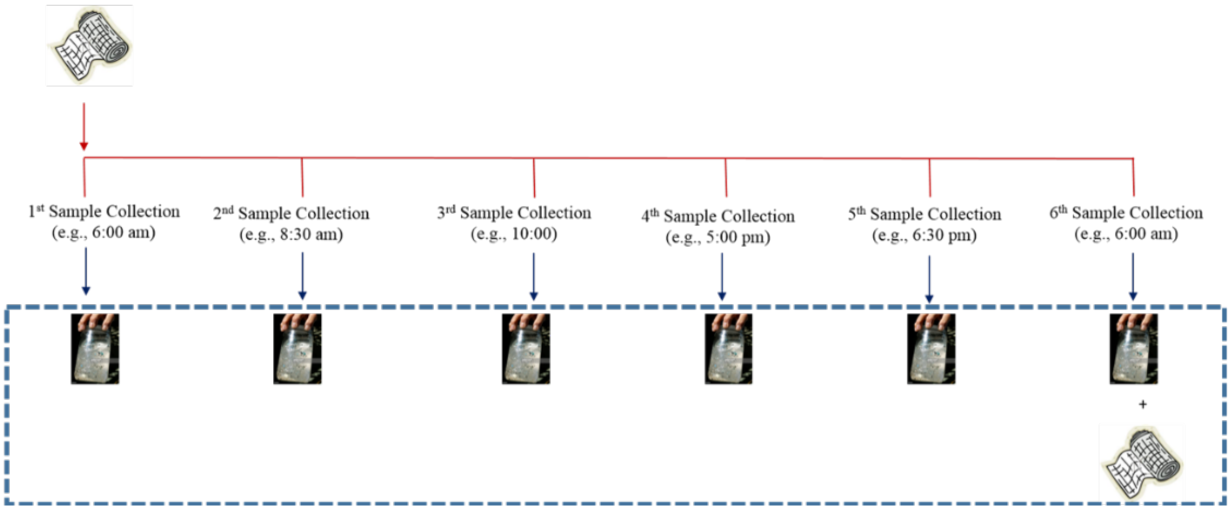

Figure S2: NY streams were enrolled by identifying watersheds with an area of ≥ 15 km$^2$ and where produce had been grown in ≥ 4 of the last 8 years for which data was available (2008-2015; 1A), and then randomly selecting six publicly accessible locations along streams in these watersheds that were ≤ 400 m from a produce field (1B).

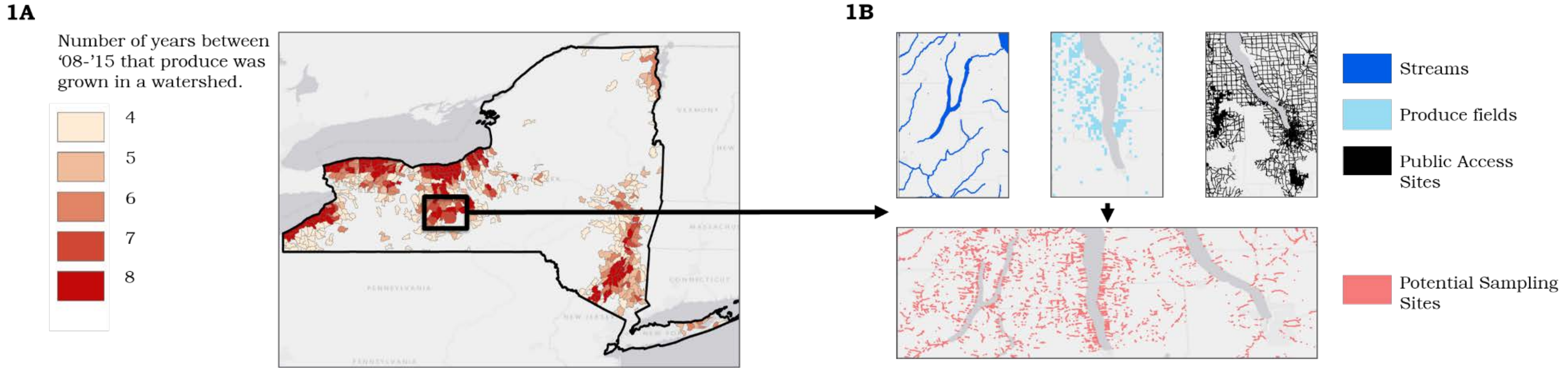

Figure S3: Scatterplots showing environmental conditions over the course of the study in Arizona. The x-axis is number of days since Jan. 1st.

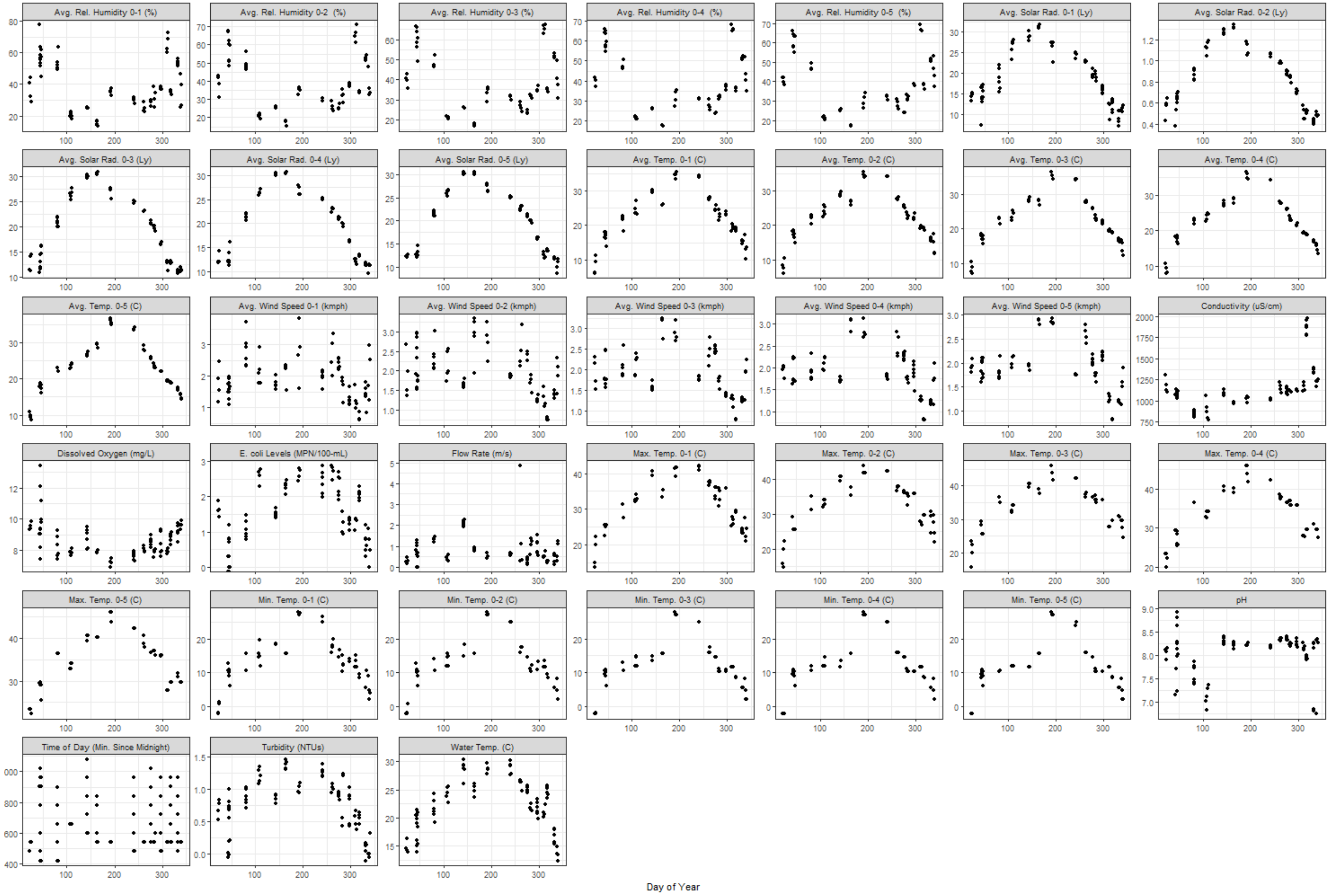

Figure S4: Scatterplots showing environmental conditions over the course of the study in New York. The x-axis is number of days since Jan. 1st.

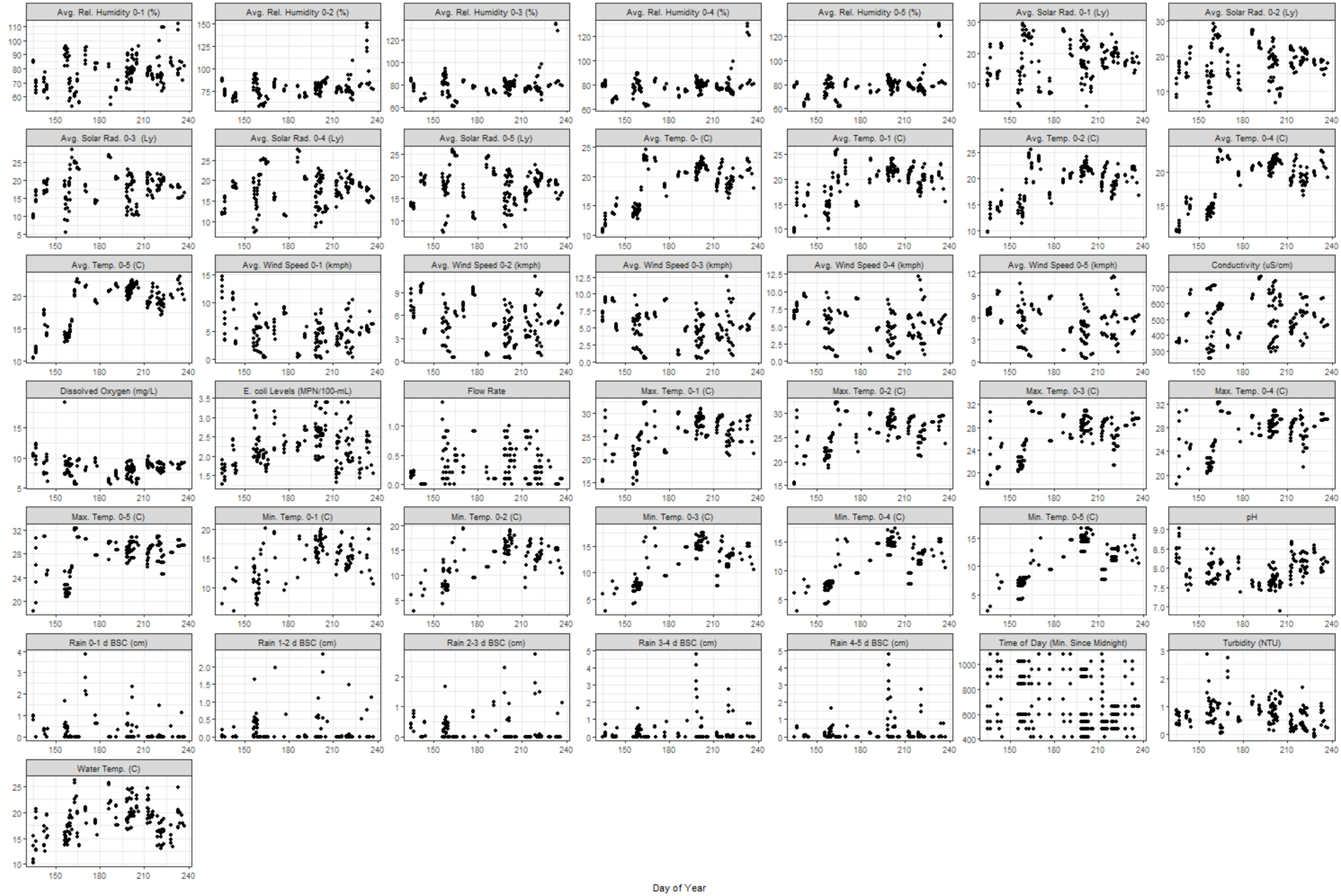

Figure S5: Matrix showing correlation between environmental factors in Arizona.

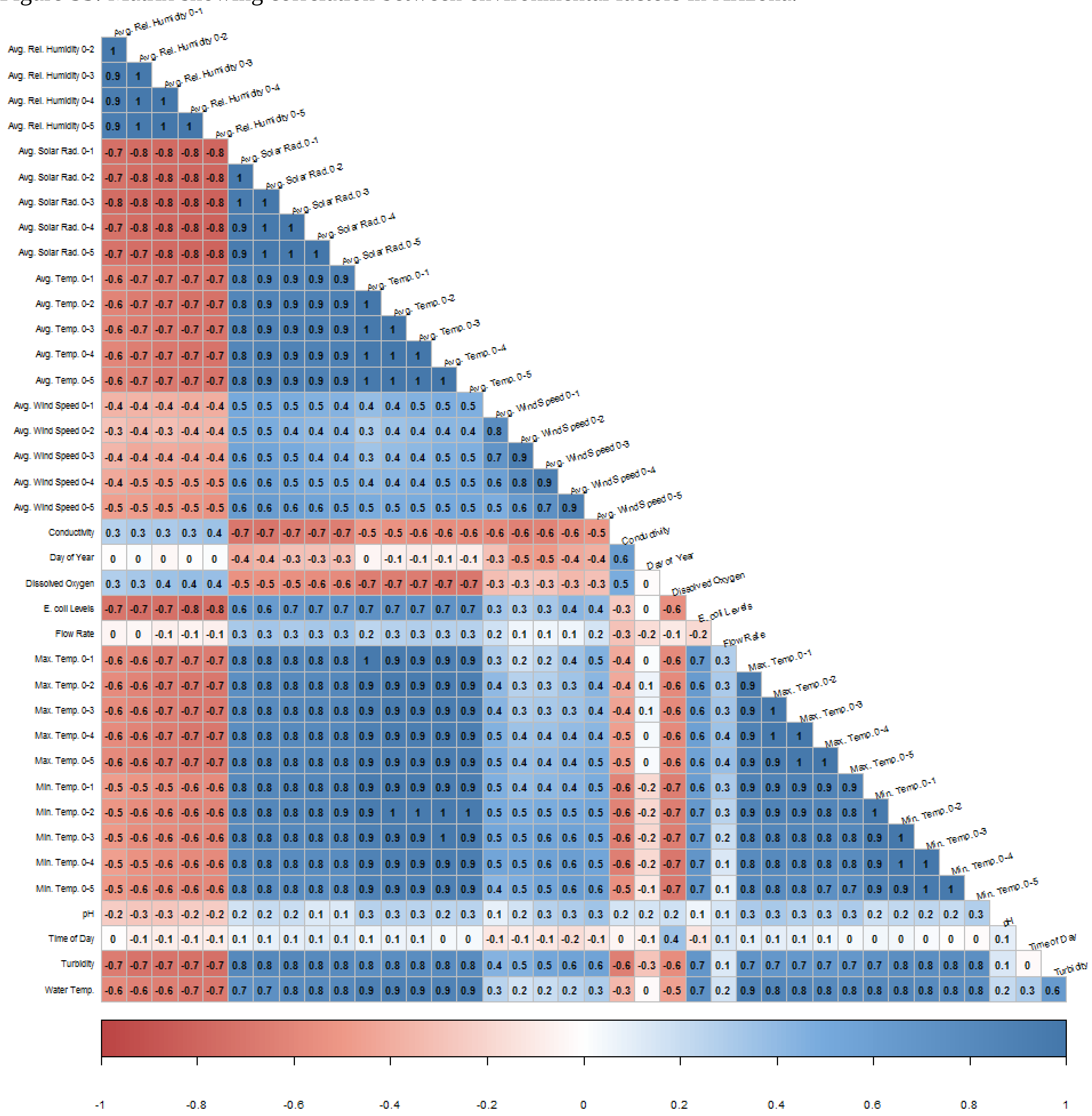

Figure S6: Matrix showing correlation between environmental factors in New York.

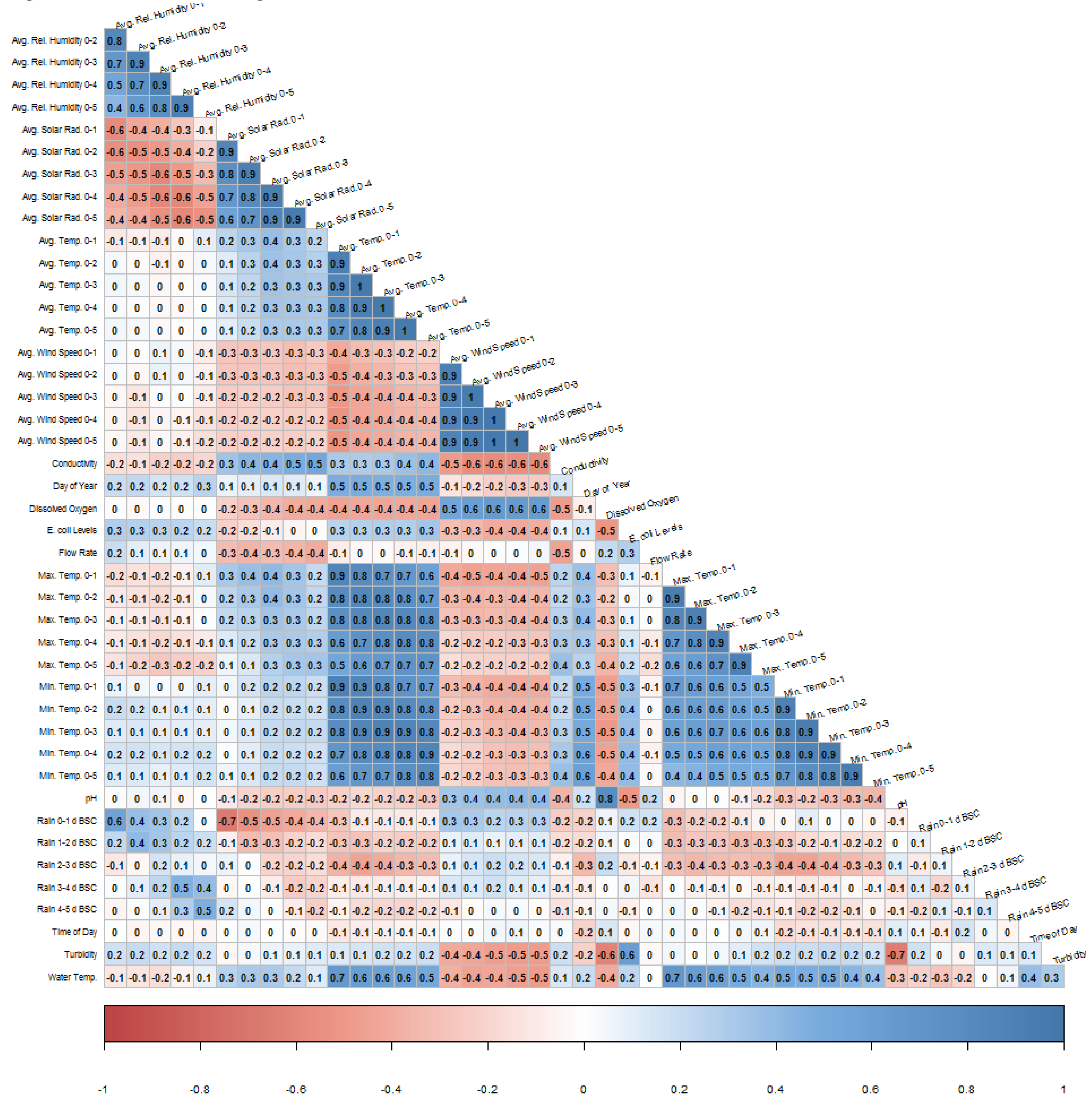

Figure S7: Partial dependence plots for the top-ranked factors associated with *E. coli* levels in Arizona or New York according to random forest analysis. The y-axis shows the factor's conditional effect on the estimated MPN of *E. coli*/100-mL, and should be interpreted as the relative impact of the outcome. For example, in AZ as dissolved oxygen increased *E. coli* levels, on average, decreased. The black lines along the x-axis show the distribution of the factor; three outliers for turbidity in NY (177, 564 and 737 NTUs) are not shown.

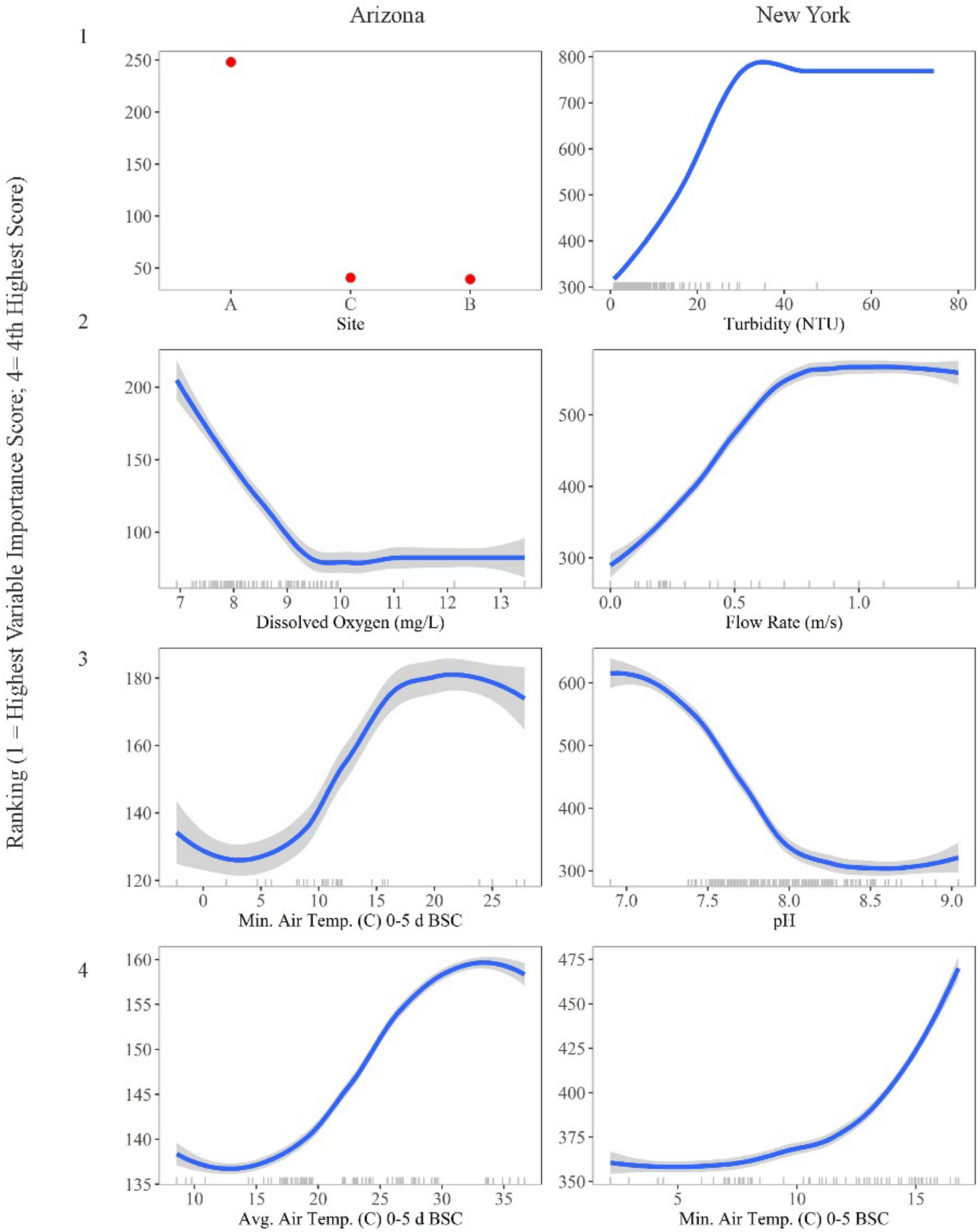

Figure S8: Results of random forest analyses that identified environmental factors associated with detecting pathogens in AZ or NY Moore swabs. The y-axis shows the factors ranked from most to least important. The x-axis shows normalized variable importance (NVI). A higher NVI (relative to the other factors) equates to a stronger association between outcome and factor; a NVI ≤ 0 indicates no association. Analyses were not performed to identify factors associated with *eaeA*-*stx* codetection or *Listeria* isolation in AZ due to the low frequency of *eaeA*-*stx*-negative and *Listeria*-positive Moore swabs in AZ. BSC = before sample collection.

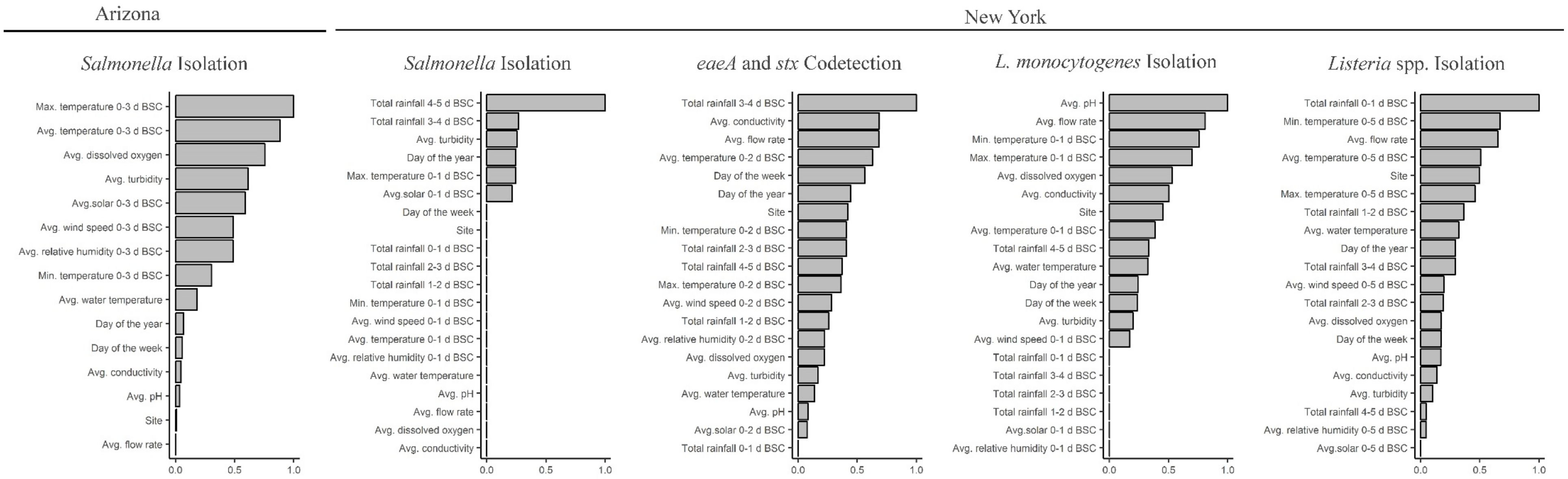

Figure S9: Partial dependence plots for the top-ranked factors associated with *L. monocytogenes* or *Listeria* spp. isolation in NY according to random forest analysis. The y-axis shows each factor's conditional effect on likelihood of isolation.

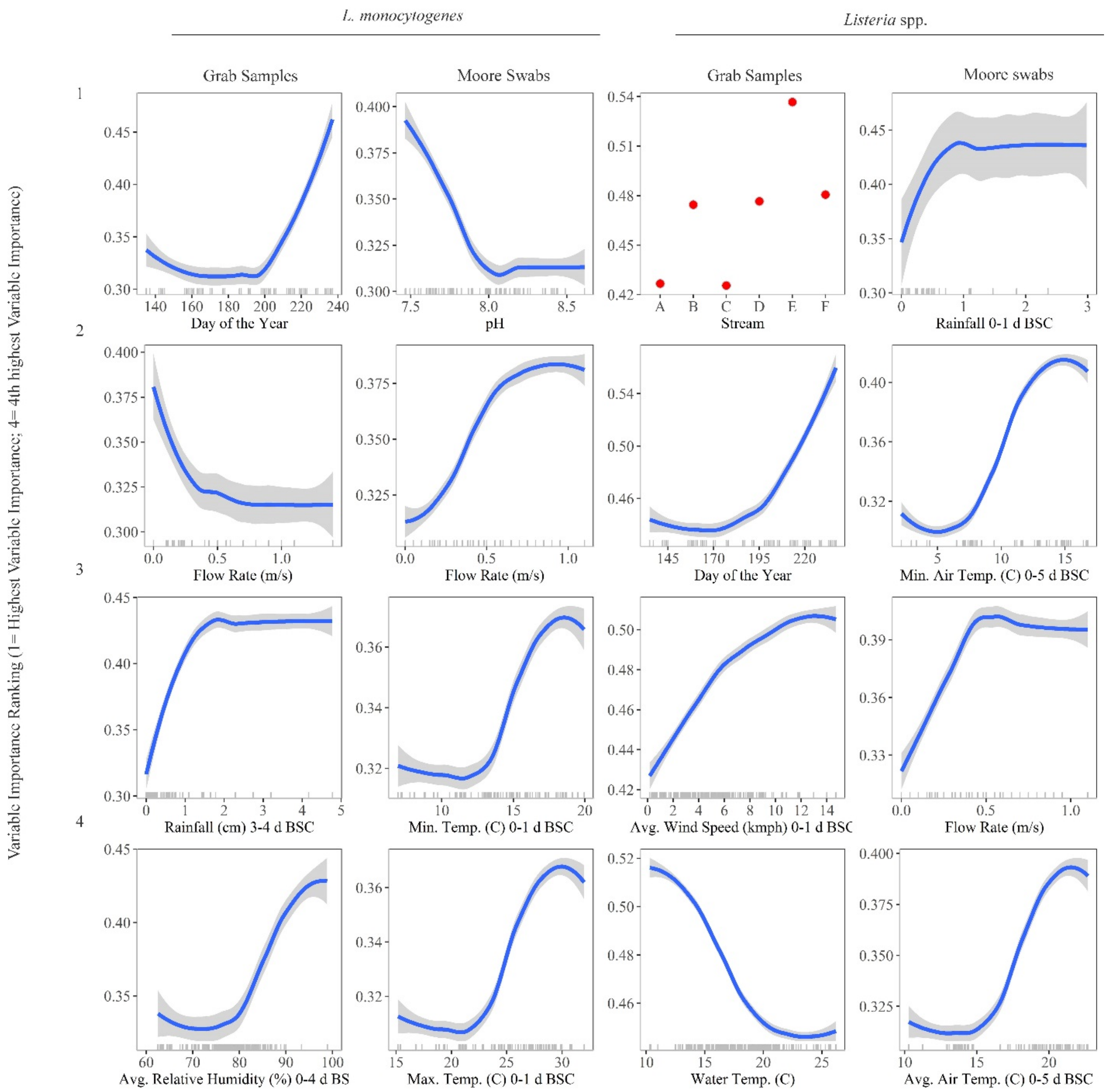

Figure S10: Partial dependence plots for the top-ranked factors associated with *Salmonella* isolation in AZ and NY according to random forest analysis. The y-axis shows each factor's conditional effect on likelihood of isolation; three outliers for turbidity in NY (at 177, 564 and 737 NTUs) are not shown.

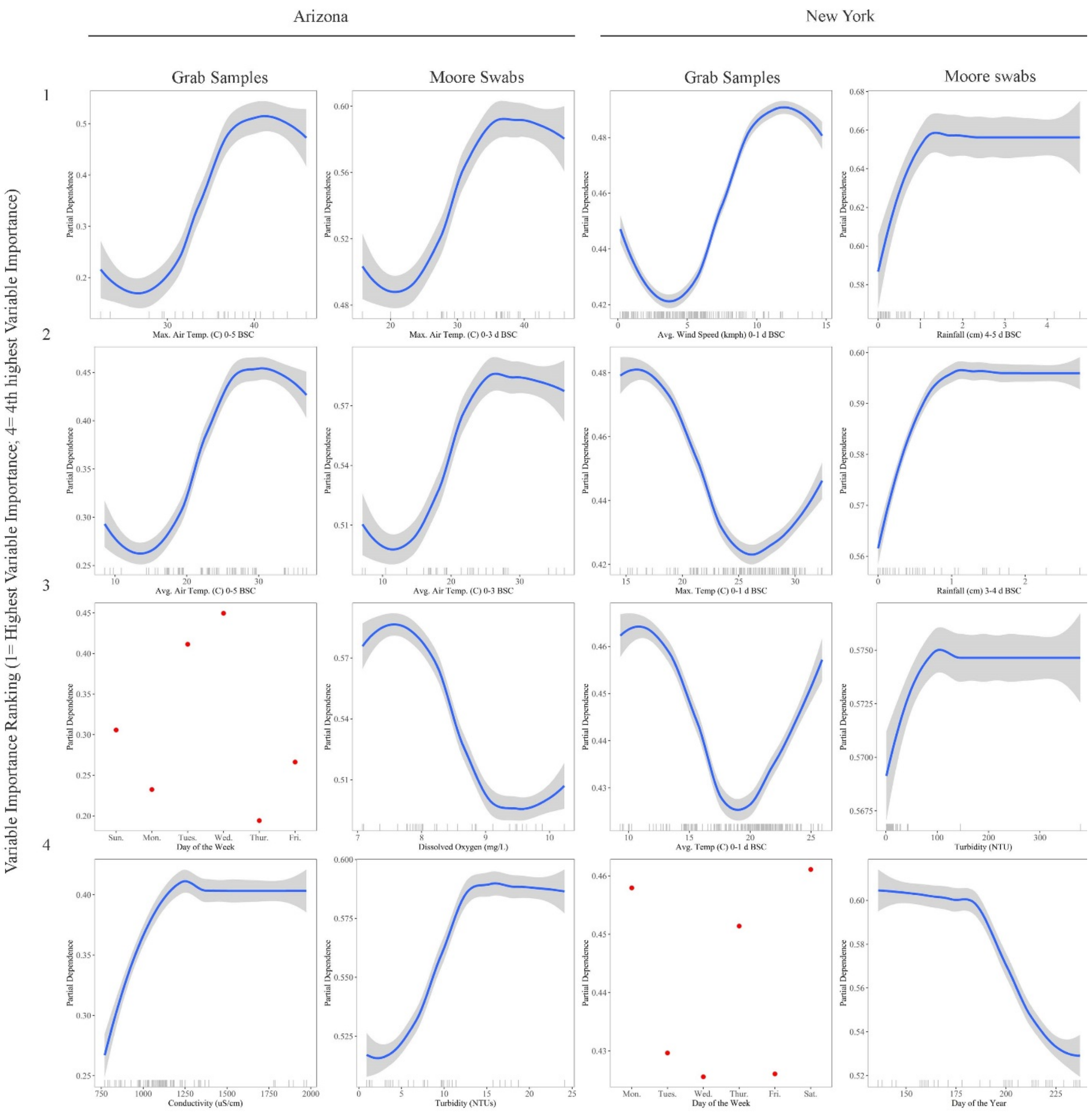

Figure S11: Partial dependence plots for the four top-ranked factors associated with codetecting *eaeA* and *stx* in AZ and NY according to random forest analysis. The y-axis shows each factor's conditional effect on likelihood of codetection.

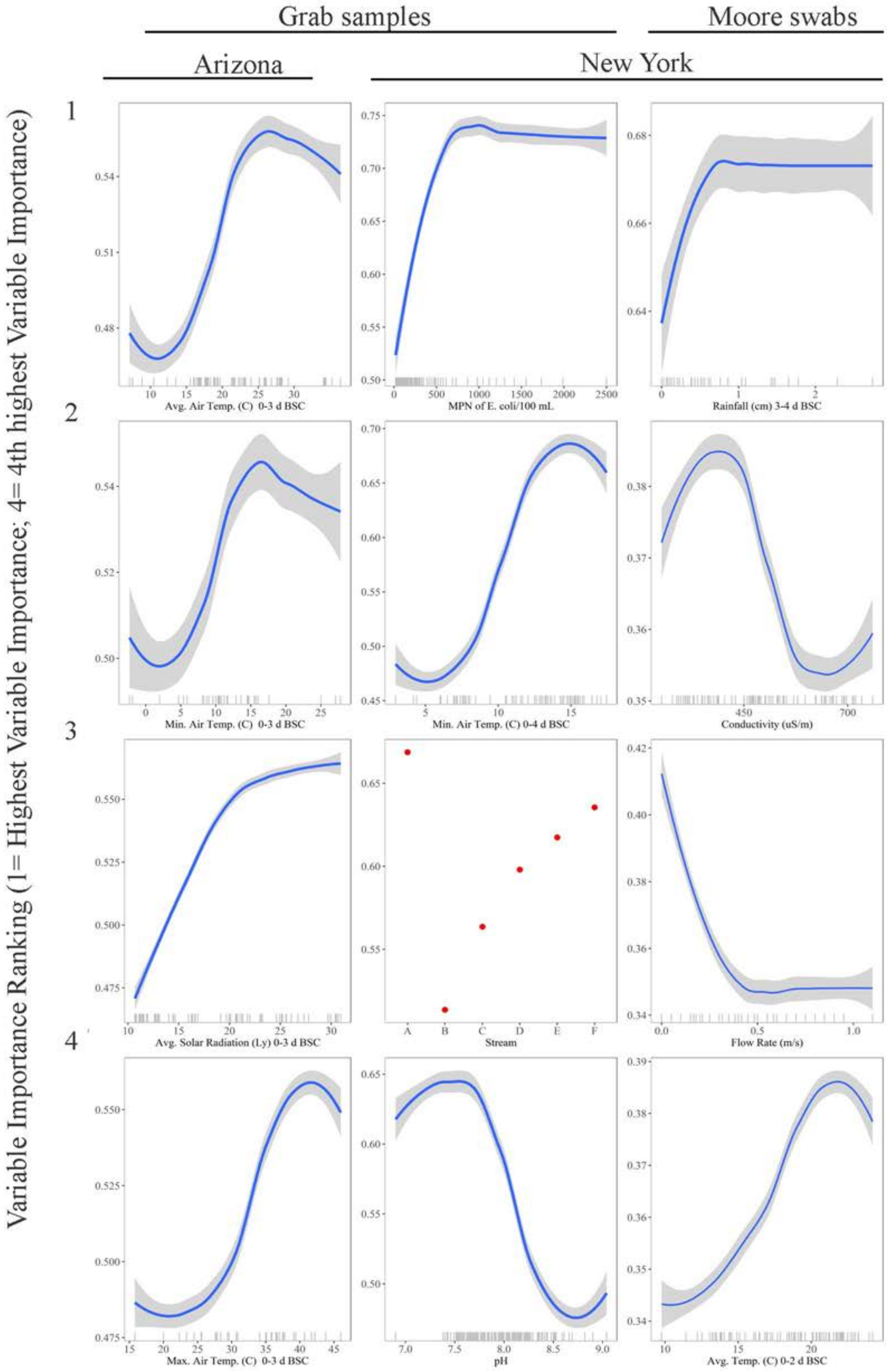

Figure S12: Partial dependence plots that illustrate two-way interactions between dissolved oxygen (DO) and other environmental factors according to random forest analyses where estimated *E. coli* levels or likelihood of pathogen detection in AZ grab samples was the outcome. Plot color ranges from white (low estimated MPN of *E. coli*/100-mL; low likelihood of pathogen detection) to dark red (high estimated MPN of *E. coli*/100-mL; high likelihood of pathogen detection). The plot that depicts the effect of pH and DO on *E. coli* levels in AZ is representative of a graph where no interaction is present since the effect of DO is constant across all levels of pH. A blank graph means that the factor on the y-axis was not associated with the outcome; for example, water temperature was not associated with *E. coli* levels in AZ.

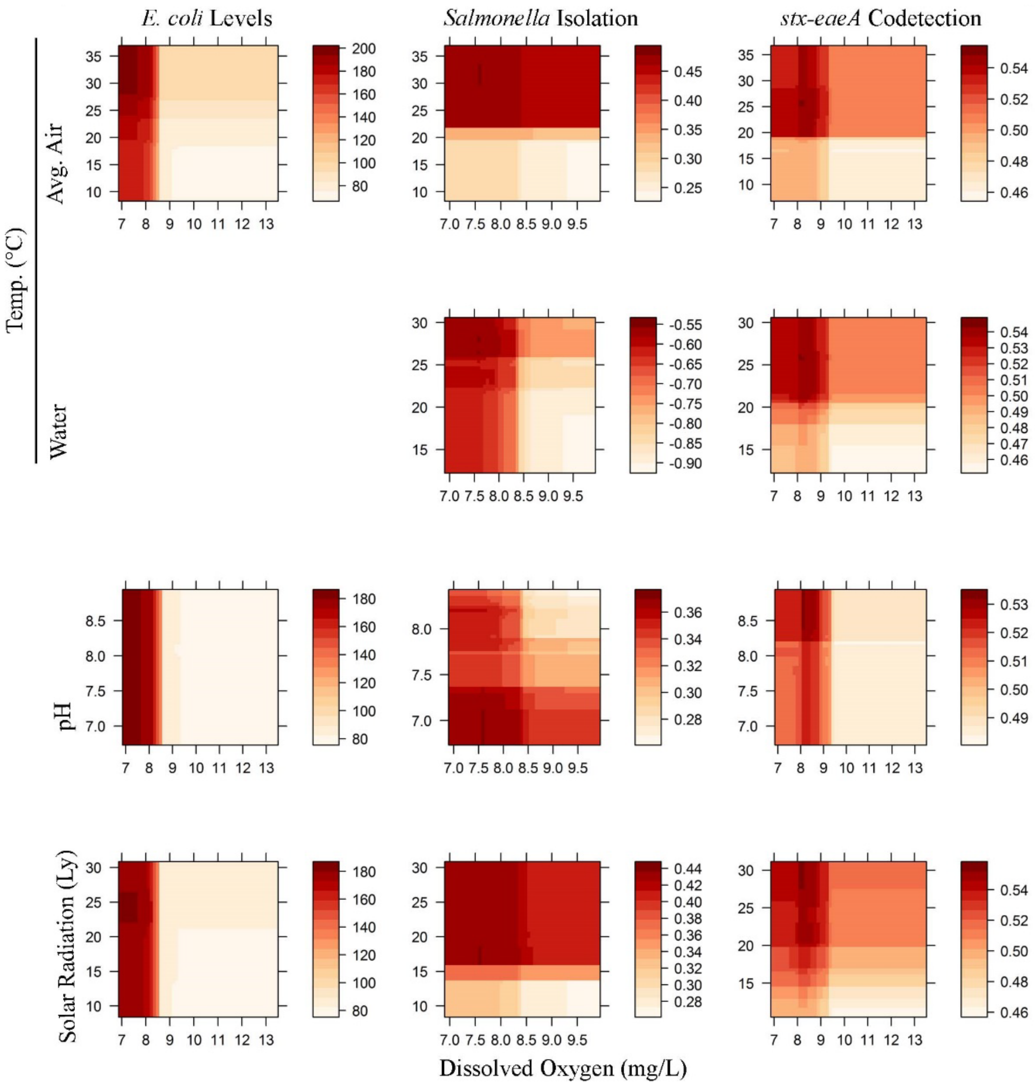

Figure S13: Partial dependence plots that illustrate two-way interactions between dissolved oxygen and other environmental factors according to random forest analyses where estimated *E. coli* levels or likelihood of pathogen detection in NY grab samples was the outcome. Plot color ranges from white (low estimated MPN of *E. coli*/100-mL; low likelihood of pathogen detection) to dark red (high estimated MPN of *E. coli*/100-mL; high likelihood of pathogen detection). A blank graph means that the factor on the y-axis was not associated with the outcome.

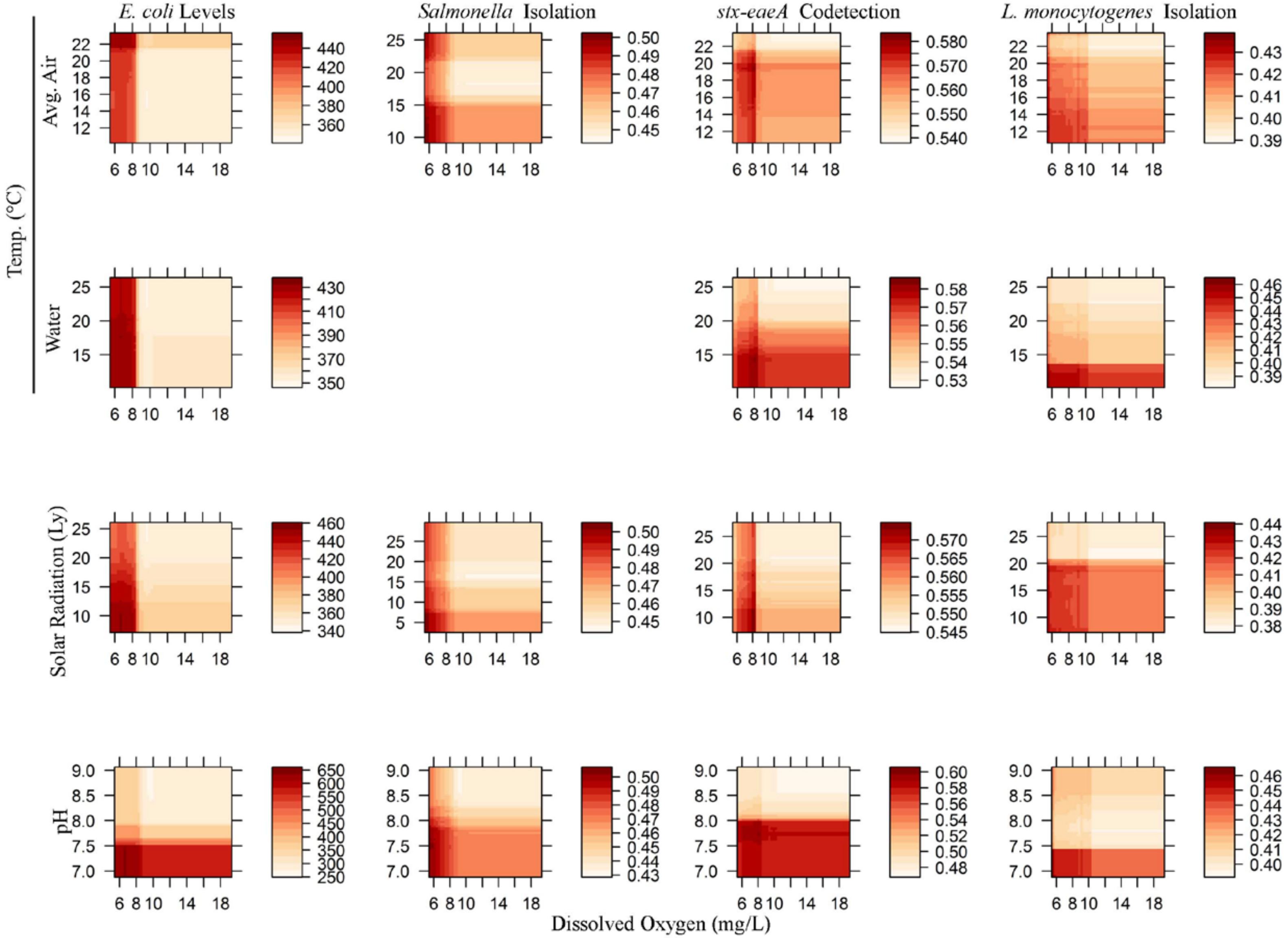

Figure S14: Partial dependence plots that illustrate two-way interactions between turbidity and other environmental factors according to random forest analyses where estimated *E. coli* levels or likelihood of pathogen detection in AZ grab samples was the outcome. Plot color ranges from white (low estimated MPN of *E. coli*/100-mL; low likelihood of pathogen detection) to dark red (high estimated MPN of *E. coli*/100-mL; high likelihood of pathogen detection). A blank graph means that the factor on the y-axis was not associated with the given outcome.

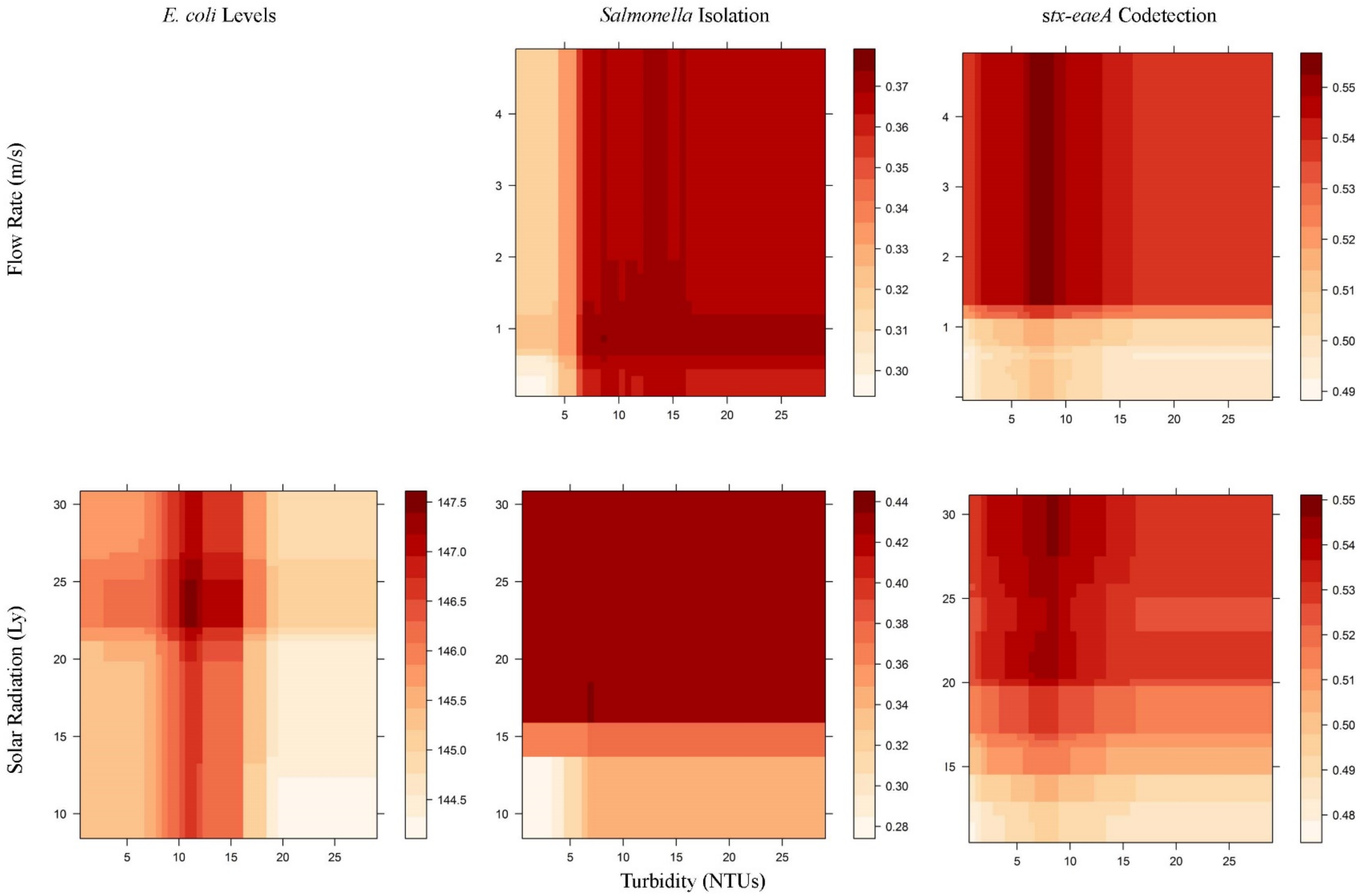

Figure S15: Partial dependence plots that illustrate two-way interactions between turbidity and other environmental factors according to random forest analyses where estimated *E. coli* levels or likelihood of pathogen detection in NY grab samples was the outcome. Plot color ranges from white (low estimated MPN of *E. coli*/100-mL; low likelihood of pathogen detection) to dark red (high estimated MPN of *E. coli*/100-mL; high likelihood of pathogen detection). A blank graph means that the factor on the y-axis was not associated with the given outcome.

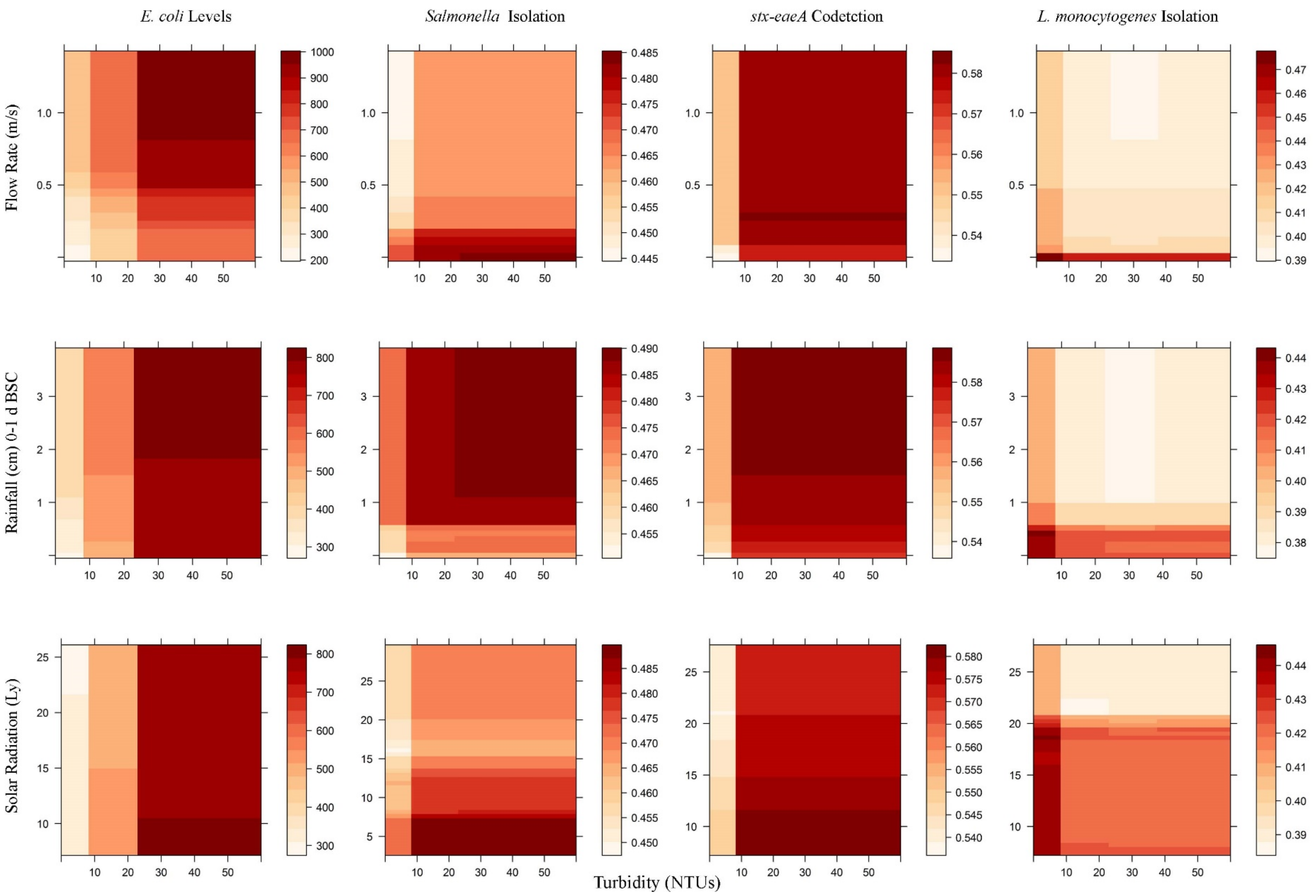

Figure S16: Partial dependence plots that illustrate two-way interactions between *E. coli* levels and physiochemical water quality factors according to random forest analyses where likelihood of pathogen detection in AZ grab samples was the outcome. Plot color ranges from white (low likelihood of pathogen detection) to dark red (high likelihood of pathogen detection).

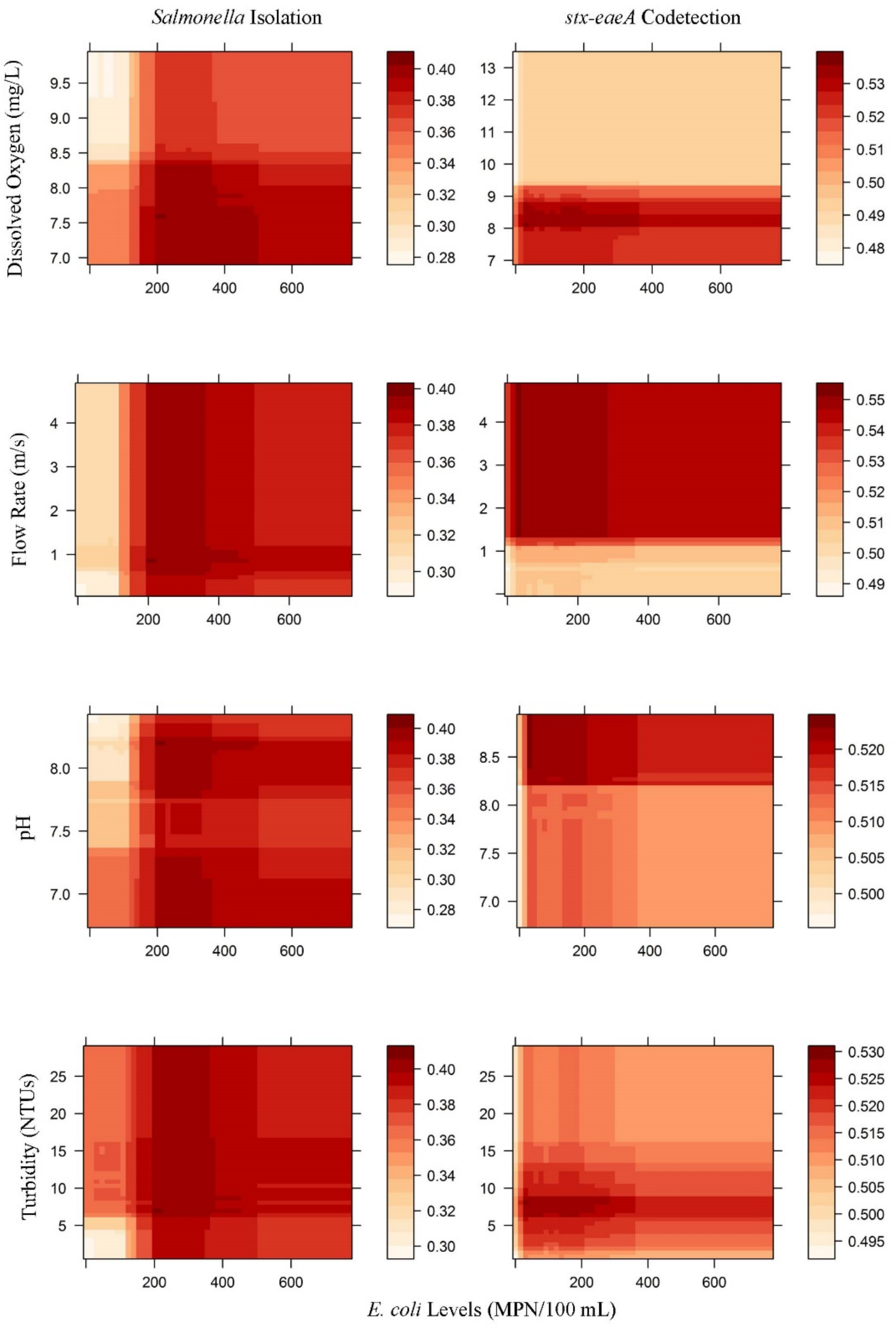

Figure S17: Partial dependence plots that illustrate two-way interactions between *E. coli* levels and weather factors according to random forest analyses where likelihood of pathogen detection in AZ grab samples was the outcome. Plot color ranges from white (low likelihood of pathogen detection) to dark red (high likelihood of pathogen detection).

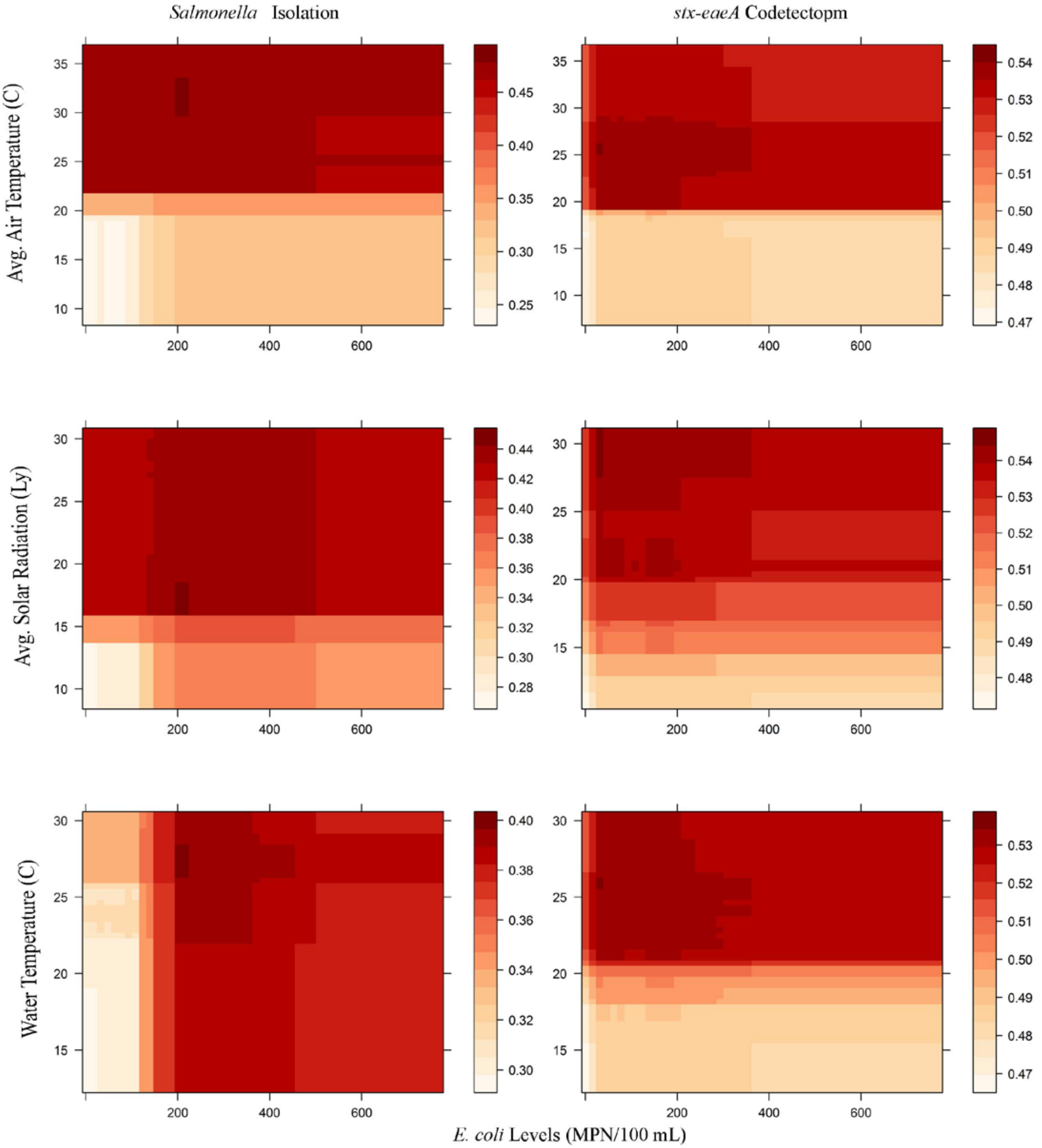

Figure S18: Partial dependence plots that illustrate two-way interactions between *E. coli* levels and physiochemical water quality factors according to random forest analyses where likelihood of pathogen detection in NY grab samples was the outcome. Plot color ranges from white (low likelihood of pathogen detection) to dark red (high likelihood of pathogen detection).

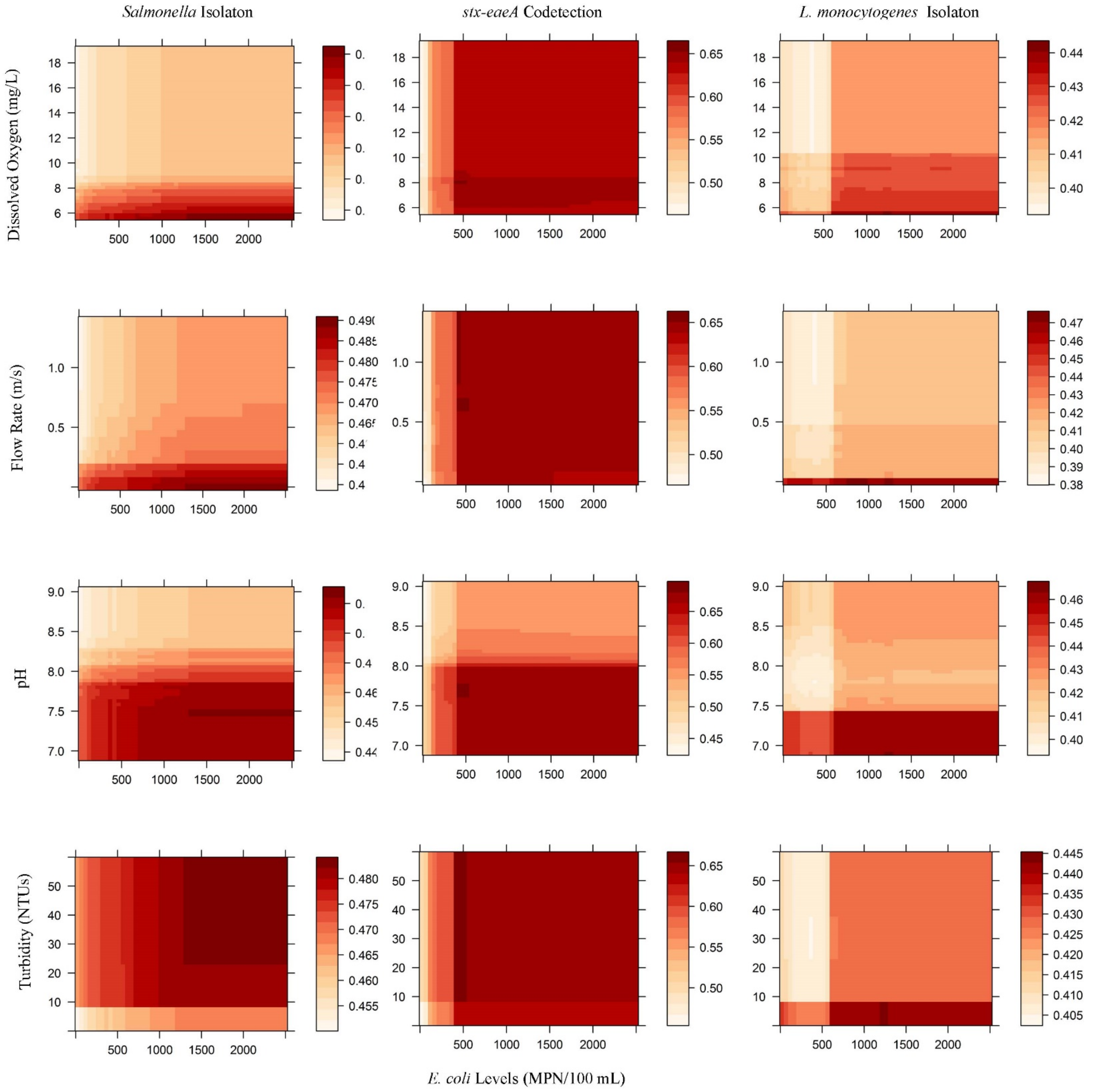

Figure S19: Partial dependence plots that illustrate two-way interactions between *E. coli* levels and weather factors according to random forest analyses where likelihood of pathogen detection in NY grab samples was the outcome. Plot color ranges from white (low likelihood of pathogen detection) to dark red (high likelihood of pathogen detection).

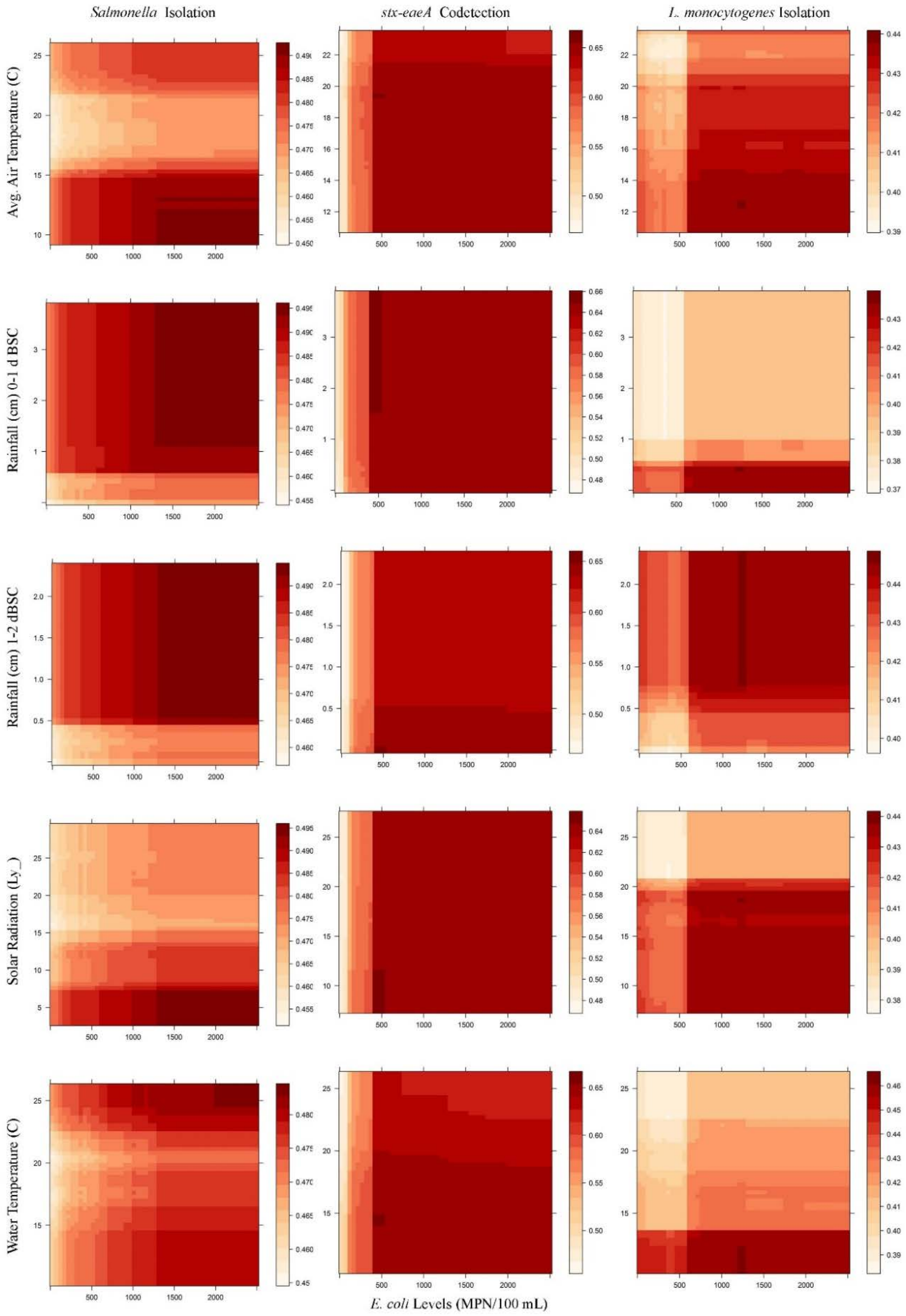

Table S1: Factors included in random forest analyses where estimated *E. coli* levels or likelihood of pathogen detection in grab samples was the outcome; rainfall factors were not included when analyzing the AZ data since no rain fell during the time periods considered in the study reported here. Values for all weather factors with the exception of rainfall were calculated for 0-1, 0-2, 0-3, 0-4, and 0-5 d before sample collection (BSC).

| Factor | Description |
|---|---|
| Avg. relative humidity | Average relative humidity 9%) |
| Avg. solar radiation | Average solar radiation (Ly) |
| Avg. air temperature | Average temperature (°C) |
| Avg. wind speed | Average wind speed (km/h) |
| Conductivity | Conductivity at time of sample collection (uS/cm) |
| Day of the week | Day of the week that the sample was collected on |
| Year-day | No. of days since Jan. 1st (i.e., Jan. 1st is year day 0, Jan. 2nd is year day 1) |
| Dissolved oxygen (DO) | Dissolved oxygen levels at time of sample collection (mg/L) |
| Flow rate [a] | Flow rate at time of sample collection (m/s) |
| Max. air temperature | Maximum temperature (°C) |
| Min. air temperature | Minimum temperature (°C) |
| *E. coli* Levels | *E. coli* concentration in the waterway (MPN of *E. coli*/100 mL) |
| pH | pH at time of sample collection |
| Site | Waterway that was sampled (i.e., Canal A, B or C in AZ, or Stream A, B, C, D, E or F in NY) |
| Time of day | Time (min. since midnight) of sample collection |
| Total rainfall 0-1 d BSC | Total rainfall (cm) on the day of sample collection |
| Total rainfall 1-2 d BSC | Total rainfall (cm) 1 to 2 days before sample collection |
| Total rainfall 2-3 d BSC | Total rainfall (cm) 2 to 3 days before sample collection |
| Total rainfall 3-4 d BSC | Total rainfall (cm) 3 to 4 days before sample collection |
| Total rainfall 4-5 d BSC | Total rainfall (cm) 4 to 5 days before sample collection |
| Turbidity | Turbidity at time of sample collection (NTUs) |
| Water temperature | Water temperature at the time of sample collection (°C) |

[a] Flow rate in AZ was measured at the surface while flow rate in NY was measured 3-6” below the surface see Methods for more details.

Table S2: Factors included in random forest analyses where likelihood of pathogen detection in Moore swabs was the outcome; rainfall factors were not included when analyzing the AZ data since no rain fell during the time periods considered in the study reported here. Values for all weather factors with the exception of rainfall were calculated for 0-1, 0-2, 0-3, 0-4, and 0-5 d before sample collection (BSC).

| Factor | Description |
|---|---|
| Avg. relative humidity | Average relative humidity |
| Avg. solar radiation | Average solar radiation (Ly) |
| Avg. air temperature | Average temperature (°C) |
| Avg. wind speed | Average wind speed (km/h) |
| Conductivity | Average conductivity while the swab was in the waterway (uS/cm) |
| Day of the week | Week day that the sample was collected on |
| Year-day | No. of days since Jan. 1$^{st}$ (i.e., Jan. 1$^{st}$ is year day 0, Jan. 2$^{nd}$ is year day 1) |
| Dissolved oxygen (DO) | Average dissolved oxygen levels while the swab was in the waterway (mg/L) |
| Flow rate [a] | Average flow rate while the swab was in the waterway (m/s) |
| Max. air temperature | Maximum temperature (°C) |
| Min. air temperature | Minimum temperature (°C) |
| pH | Average pH while the swab was in the waterway |
| Site | The waterway that was sampled (i.e., Canal A, B or C in AZ, or Stream A, B, C, D, E or F in NY) |
| Total rainfall 0-1 d BSC | Total rainfall (cm) on the day of sample collection |
| Total rainfall 1-2 d BSC | Total rainfall (cm) 1 to 2 days before sample collection |
| Total rainfall 2-3 d BSC | Total rainfall (cm) 2 to 3 days before sample collection |
| Total rainfall 3-4 d BSC | Total rainfall (cm) 3 to 4 days before sample collection |
| Total rainfall 4-5 d BSC | Total rainfall (cm) 4 to 5 days before sample collection |
| Turbidity | Average turbidity while the Moore swab was in the waterway (NTUs) |
| Water temperature | Average water temperature a while the Moore swab was in the waterway (°C) |

[a] Flow rate in AZ was measured at the surface while flow rate in NY was measured 3-6" below the surface see Methods for more details.

Table S3: Results of RFs that identified factors associated with *E. coli* levels. Five, overlapping time frames [0-1, 0-2, 0-3, 0-4, or 0-5 d before sample collection (BSC)] were used to calculate the values of weather factors with the exception of rainfall; rainfall was calculated daily (0-1, 1-2, 2-3, 3-4, 4-5 d BSC). Separate random forests were developed for each outcome and overlapping time frame in each state. Results are reported as the rank each factor received (e.g., 1st means that the factor was the top-ranked factor); U = uninformative.

| | Arizona | | | | | New York | | | | |
|---|---|---|---|---|---|---|---|---|---|---|
| Factors | 0-1 | 0-2 | 0-3 | 0-4 | 0-5 | 0-1 | 0-2 | 0-3 | 0-4 | 0-5 |
| Avg. Air Temp. | 3rd | 3rd | 3rd | 4th | 4th | 16th | 14th | 12th | 9th | 12th |
| Avg. Relative Humidity | 8th | 8th | 8th | U | 7th | 6th | 12th | 11th | 14th | 11th |
| Avg. Solar Radiation | 7th | U | 7th | 9th | 9th | 4th | 5th | 8th | 10th | 8th |
| Avg. Wind Speed | U | U | U | U | U | 12th | 6th | 4th | 5th | 4th |
| Conductivity | 9th | U | 9th | 8th | 8th | 13th | 13th | 15th | 13th | 14th |
| Day of the Week | U | U | U | U | U | U | U | U | U | U |
| Year-day | 11th | 10th | U | 12th | 10th | 7th | 9th | 9th | 11th | 10th |
| Dissolved Oxygen | 2nd | 2nd | 2nd | 2nd | 2nd | 5th | 4th | 5th | 6th | 6th |
| Flow Rate | U | 7th | U | 7th | U | 2nd | 2nd | 2nd | 2nd | 2nd |
| Max. Air Temp. | 4th | 9th | 10th | 11th | U | 11th | 11th | 13th | 17th | 16th |
| Min. Air Temp. | U | 5th | 4th | 3rd | 3rd | 8th | 7th | 6th | 4th | 5th |
| pH | U | U | U | U | U | 3rd | 3rd | 3rd | 3rd | 3rd |
| Site | 1st | 1st | 1st | 1st | 1st | 9th | 10th | 10th | 8th | 9th |
| Time Since Midnight | 5th | 4th | 5th | 5th | 5th | 18th | 18th | 20th | U | U |
| Rainfall 0-1 d BSC | NA | NA | NA | NA | NA | 10th | 8th | 7th | 7th | 7th |
| Rainfall 1-2 d BSC | NA | NA | NA | NA | NA | 14th | U | 17th | 18th | 17th |
| Rainfall 2-3 d BSC | NA | NA | NA | NA | NA | 17th | U | 19th | U | 18th |
| Rainfall 3-4 d BSC | NA | NA | NA | NA | NA | 19th | 16th | 16th | 16th | U |
| Rainfall 4-5 d BSC | NA | NA | NA | NA | NA | U | 17th | 18th | 15th | 15th |
| Turbidity | 6th | 6th | 6th | 6th | 6th | 1st | 1st | 1st | 1st | 1st |
| Water Temp. | 10th | U | U | 10th | U | 15th | 15th | 14th | 12th | 13th |
| $R^2$ | 0.71 | 0.71 | 0.71 | 0.71 | 0.72 | 0.43 | 0.44 | 0.44 | 0.45 | 0.45 |

Table S4: Results of RFs that identified factors associated with *L. monocytogenes* or *Listeria* spp. isolation in NY. Five, overlapping time frames [0-1, 0-2, 0-3, 0-4, or 0-5 d before sample collection (BSC)] were used to calculate the values of weather factors with the exception of rainfall; rainfall was calculated daily (0-1, 1-2, 2-3, 3-4, 4-5 d BSC). Separate random forests were developed for each outcome and overlapping time frame in each state. Results are reported as the rank each factor received (e.g., 1st means that the factor was the top-ranked factor); U = uninformative.

| Factors | *L. monocytogenes* | | | | | | | | | | *Listeria* spp. | | | | | | | | | |
|---|---|---|---|---|---|---|---|---|---|---|---|---|---|---|---|---|---|---|---|---|
| | Grab samples | | | | | Moore swabs | | | | | Grab samples | | | | | Moore swabs | | | | |
| | 0-1 | 0-2 | 0-3 | 0-4 | 0-5 | 0-1 | 0-2 | 0-3 | 0-4 | 0-5 | 0-1 | 0-2 | 0-3 | 0-4 | 0-5 | 0-1 | 0-2 | 0-3 | 0-4 | 0-5 |
| Avg. Air Temp. | 12th | 15th | 14th | 12th | 16th | 8th | 4th | 4th | 5th | 7th | 19th | 17th | U | 14th | 15th | 8th | 3rd | 9th | 8th | 4th |
| Avg. Relative Humidity | 14th | 6th | 2nd | 4th | 4th | U | 14th | U | 16th | 1st | 14th | 9th | 8th | 9th | 3rd | 19th | 10th | 19th | 19th | 19th |
| Avg. Solar Radiation | 7th | 5th | 13th | 7th | 3rd | U | 12th | U | 12th | 5th | 5th | 10th | 17th | U | 16th | 12th | 17th | 20th | 20th | 20th |
| Avg. Wind Speed | 9th | 14th | 9th | 8th | 9th | 14th | 17th | 12th | 15th | 17th | 3rd | 12th | 12th | 13th | 7th | 20th | 20th | 18th | 12th | 11th |
| Conductivity | 15th | 17th | 17th | 17th | 17th | 6th | 6th | 6th | 6th | 6th | 12th | 11th | 6th | 7th | 10th | 11th | 15th | 13th | 14th | 16th |
| Day of the Week | 4th | 7th | 8th | 9th | 8th | 12th | 13th | 9th | 11th | 15th | U | U | U | U | U | 13th | 12th | 8th | 6th | 14th |
| Year-day | 2nd | 2nd | 3rd | 1st | 1st | 11th | 10th | 10th | 8th | 13th | 2nd | 2nd | 2nd | 2nd | 2nd | 5th | 6th | 6th | 7th | 9th |
| Dissolved Oxygen | 8th | 9th | 7th | 21st | 11th | 5th | 7th | 5th | 4th | 10th | 10th | 14th | 15th | 15th | 11th | 14th | 14th | 15th | 15th | 13th |
| E. coli Levels | 16th | 19th | 21st | 20th | 19th | NA | NA | NA | NA | NA | 17th | U | 13th | U | 17th | NA | NA | NA | NA | NA |
| Flow Rate | 1st | 1st | 1st | 2nd | 2nd | 2nd | 2nd | 2nd | 2nd | 4th | 9th | 4th | 4th | 4th | 9th | 3rd | 4th | 3rd | 3rd | 3rd |
| Max. Air Temp. | 13th | 4th | 12th | 19th | 18th | 4th | 3rd | 11th | 13th | 12th | 15th | 7th | 16th | U | 18th | 10th | 11th | 11th | 10th | 6th |
| Min. Air Temp. | 19th | 18th | 16th | 11th | 5th | 3rd | 8th | 3rd | 3rd | 3rd | 18th | 15th | U | 10th | 5th | 2nd | 2nd | 2nd | 2nd | 2nd |
| pH | 5th | 3rd | 5th | 6th | 6th | 1st | 1st | 1st | 1st | 2nd | 6th | 3rd | 3rd | 3rd | 4th | 15th | 19th | 14th | 17th | 15th |
| Site | 6th | 8th | 11th | 10th | 10th | 7th | 9th | 8th | 7th | 8th | 1st | 1st | 1st | 1st | 1st | 4th | 8th | 4th | 4th | 5th |
| Min. Since Midnight | 18th | 16th | 19th | 18th | 15th | NA | NA | NA | NA | NA | U | U | U | U | U | NA | NA | NA | NA | NA |
| Rainfall 0-1 d BSC | 22nd | 20th | 18th | 15th | 21st | U | 16th | U | U | 16th | 8th | 6th | 5th | 5th | 8th | 1st | 1st | 1st | 1st | 1st |
| Rainfall 1-2 d BSC | 21st | 21st | 20th | 14th | 20th | U | U | U | U | U | U | U | U | U | U | 7th | 5th | 7th | 5th | 7th |
| Rainfall 2-3 d BSC | 20th | 22nd | 22nd | 22nd | 22nd | U | 18th | U | U | U | 11th | 13th | 10th | 12th | 12th | 16th | 13th | 12th | 16th | 12th |
| Rainfall 3-4 d BSC | 3rd | 12th | 10th | 3rd | 12th | U | U | U | U | U | 16th | 16th | 11th | 11th | 13th | 6th | 7th | 5th | 11th | 1 0th |
| Rainfall 4-5 d BSC | 10th | 11th | 4th | 13th | 7th | 9th | 5th | 14th | 10th | 9th | 7th | 8th | 9th | 8th | 14th | 17th | 18th | 16th | 13th | 18th |
| Turbidity | 17th | 10th | 6th | 5th | 13th | 13th | 15th | 13th | 14th | 14th | 13th | U | 14th | U | U | 18th | 16th | 17th | 18th | 17th |
| Water Temp. | 11th | 13th | 15th | 16th | 14th | 10th | 11th | 7th | 9th | 11th | 4th | 5th | 7th | 6th | 6th | 9th | 9th | 10th | 9th | 8th |

| | | | | | | | | | | | | | | | | | | | | |
|---|---|---|---|---|---|---|---|---|---|---|---|---|---|---|---|---|---|---|---|---|
| Accuracy | 0.765 | 0.752 | 0.758 | 0.757 | 0.747 | 0.900 | 0.897 | 0.901 | 0.897 | 0.900 | 0.686 | 0.672 | 0.669 | 0.669 | 0.674 | 0.731 | 0.740 | 0.720 | 0.740 | 0.751 |
| Kappa Statistic | 0.046 | 0.028 | 0.046 | 0.060 | 0.027 | 0.366 | 0.316 | 0.357 | 0.323 | 0.345 | 0.363 | 0.335 | 0.328 | 0.329 | 0.337 | 0.301 | 0.328 | 0.234 | 0.330 | 0.353 |

Table S5: Results of RFs that identified factors associated with *Salmonella* isolation in AZ and NY. Five, overlapping time frames [0-1, 0-2, 0-3, 0-4, or 0-5 d before sample collection (BSC)] were used to calculate the values of weather factors with the exception of rainfall; rainfall was calculated daily (0-1, 1-2, 2-3, 3-4, 4-5 d BSC). Separate random forests were developed for each outcome and overlapping time frame in each state. Results are reported as the rank each factor received (e.g., 1st means that the factor was the top-ranked factor); U = uninformative.

| | Arizona | | | | | | | | | | New York | | | | | | | | | |
|---|---|---|---|---|---|---|---|---|---|---|---|---|---|---|---|---|---|---|---|---|
| | Grab Samples | | | | | Moore Swabs | | | | | Grab Samples | | | | | Moore Swabs | | | | |
| Factors | 0-1 | 0-2 | 0-3 | 0-4 | 0-5 | 0-1 | 0-2 | 0-3 | 0-4 | 0-5 | 0-1 | 0-2 | 0-3 | 0-4 | 0-5 | 0-1 | 0-2 | 0-3 | 0-4 | 0-5 |
| Avg. Air Temp. | 11th | 3rd | 3rd | 13th | 3rd | 1st | 2nd | 2nd | 2nd | 2nd | 3rd | 13th | U | 20th | 19th | U | U | U | U | U |
| Avg. Relative Humidity | 7th | 13th | 13th | 10th | 13th | 8th | 5th | 7th | 7th | 6th | 17th | 7th | 8th | 9th | 4th | U | U | U | U | U |
| Avg. Solar Radiation | 14th | 4th | 5th | 15th | 6th | 7th | 7th | 5th | 4th | 5th | 8th | 11th | 7th | 5th | 5th | 6th | U | U | U | U |
| Avg. Wind Speed | 10th | 16th | 8th | 11th | 9th | 5th | 6th | 6th | 9th | 9th | 1st | 1st | 9th | 7th | 7th | U | U | U | U | U |
| Conductivity | 6th | 6th | 4th | 5th | 4th | 10th | 11th | 12th | 10th | 10th | U | 19th | U | 18th | 13th | U | U | U | U | U |
| Day of the Week | 2nd | 2nd | 2nd | 2nd | 2nd | 12th | 13th | 11th | 12th | 13th | 4th | 3rd | 2nd | 2nd | 2nd | U | U | U | 4th | 4th |
| Year-day | 3rd | 5th | 6th | 3rd | 5th | 11th | 10th | 10th | 11th | 11th | 15th | 16th | 12th | 10th | 11th | 4th | U | U | U | 3rd |
| Dissolved Oxygen | 5th | 8th | 9th | 6th | 10th | 2nd | 3rd | 3rd | 3rd | 3rd | 7th | 9th | 11th | 12th | 10th | U | U | U | U | U |
| E. coli Levels | 13th | 12th | 14th | 14th | 14th | NA | NA | NA | NA | NA | 19th | 20th | 14th | 19th | 17th | NA | NA | NA | NA | NA |
| Flow Rate | 9th | 10th | 12th | 8th | 12th | 15th | 15th | 15th | U | 15th | 11th | 5th | 5th | 6th | 8th | U | U | U | U | U |
| Max. Air Temp. | 1st | 1st | 1st | 1st | 1st | 3rd | 1st | 1st | 1st | 1st | 2nd | 8th | 6th | 11th | 20th | 5th | U | 2nd | 2nd | 5th |
| Min. Air Temp. | 16th | 11th | 11th | 16th | 11th | 6th | 9th | 8th | 6th | 7th | 12th | 14th | 17th | 14th | 12th | U | U | 4th | U | U |
| Min. Since Midnight | 4th | 9th | 7th | 4th | 7th | NA | NA | NA | NA | NA | U | U | U | U | U | NA | NA | NA | NA | NA |
| pH | 8th | 7th | 10th | 9th | 8th | 13th | 12th | 13th | 14th | 12th | 13th | 10th | 4th | 4th | 3rd | U | U | U | U | U |
| Site | 17th | 17th | 17th | 17th | 17th | 14th | 14th | 14th | 13th | 14th | 10th | 12th | 15th | 17th | 18th | U | U | U | U | U |
| Rainfall 0-1 d BSC | NA | NA | NA | NA | NA | NA | NA | NA | NA | NA | 5th | 2nd | 1st | 1st | 1st | U | U | U | U | U |
| Rainfall 1-2 d BSC | NA | NA | NA | NA | NA | NA | NA | NA | NA | NA | 16th | 15th | U | 15th | 14th | U | U | U | U | U |
| Rainfall 2-3 d BSC | NA | NA | NA | NA | NA | NA | NA | NA | NA | NA | 18th | U | 13th | 13th | 15th | 2nd | U | U | U | 6th |
| Rainfall 3-4 d BSC | NA | NA | NA | NA | NA | NA | NA | NA | NA | NA | 14th | 17th | 16th | 16th | 16th | 1st | 1st | 1st | 1st | 1st |
| Rainfall 4-5 d BSC | NA | NA | NA | NA | NA | NA | NA | NA | NA | NA | 9th | 4th | 3rd | 3rd | 6th | U | U | U | U | U |
| Turbidity | 12th | 14th | 15th | 7th | 16th | 4th | 4th | 4th | 5th | 4th | 6th | 6th | 10th | 8th | 9th | 3rd | 2nd | 3rd | 3rd | 2nd |

| | | | | | | | | | | | | | | | | | | | | |
|---|---|---|---|---|---|---|---|---|---|---|---|---|---|---|---|---|---|---|---|---|
| Water Temp. | 15th | 15th | 16th | 12th | 15th | 9th | 8th | 9th | 8th | 8th | U | U | U | U | U | U | U | U | U | U |
| Accuracy | 0.709 | 0.705 | 0.705 | 0.708 | 0.718 | 0.822 | 0.831 | 0.839 | 0.831 | 0.831 | 0.605 | 0.600 | 0.594 | 0.594 | 0.596 | 0.579 | 0.538 | 0.561 | 0.563 | 0.539 |
| Kappa Statistic | 0.394 | 0.361 | 0.361 | 0.392 | 0.396 | 0.630 | 0.660 | 0.677 | 0.660 | 0.660 | 0.178 | 0.166 | 0.163 | 0.165 | 0.167 | 0.110 | 0.026 | 0.067 | 0.081 | 0.010 |

Table S7: Results of RFs that identified factors associated with *eaeA-stx* codetection in AZ and NY. Five, overlapping time frames [0-1, 0-2, 0-3, 0-4, or 0-5 d before sample collection (BSC)] were used to calculate the values of weather factors with the exception of rainfall; rainfall was calculated daily (0-1, 1-2, 2-3, 3-4, 4-5 d BSC). Separate random forests were developed for each outcome and overlapping time frame in each state. Results are reported as the rank each factor received (e.g., 1st means that the factor was the top-ranked factor); U = uninformative.

| Factors | Arizona | | | | | New York | | | | | | | | | |
|---|---|---|---|---|---|---|---|---|---|---|---|---|---|---|---|
| | Grab Samples | | | | | Grab Samples | | | | | Moore Swabs | | | | |
| | 0-1 | 0-2 | 0-3 | 0-4 | 0-5 | 0-1 | 0-2 | 0-3 | 0-4 | 0-5 | 0-1 | 0-2 | 0-3 | 0-4 | 0-5 |
| Avg. Relative Humidity | 15th | 8th | 6th | 7th | 8th | 17th | 12th | 11th | 8th | 14th | 15th | 14th | 7th | 19th | U |
| Avg. Solar Radiation | 6th | 4th | 3rd | 2nd | 2nd | 4th | 19th | 19th | 19th | 21st | 18th | 19th | 10th | 14th | 17th |
| Avg. Air Temp. | 2nd | 1st | 1st | 1st | 1st | 22nd | 22nd | 17th | 7th | 9th | 4th | 4th | 8th | 5th | 6th |
| Avg. Wind Speed | 9th | 9th | 15th | 14th | 11th | 18th | 10th | 14th | 14th | 11th | 6th | 12th | 13th | 3rd | 5th |
| Conductivity | 11th | 14th | 11th | 13th | 10th | 16th | 20th | 20th | 22nd | 20th | 5th | 2nd | 3rd | 4th | 3rd |
| Dissolved Oxygen | 5th | 6th | 7th | 6th | 7th | 11th | 15th | 13th | 17th | 16th | 13th | 15th | 14th | 15th | 15th |
| E. coli Levels | 10th | 13th | 14th | 11th | 13th | 1st | 1st | 1st | 1st | 1st | NA | NA | NA | NA | NA |
| Flow Rate | 7th | 7th | 8th | 8th | 6th | 12th | 14th | 15th | 13th | 15th | 8th | 3rd | 9th | 7th | 8th |
| Max. Air Temp. | 3rd | 3rd | 4th | 4th | 3rd | 21st | 17th | 7th | 16th | 17th | 3rd | 11th | 11th | 2nd | 2nd |
| Min. Air Temp. | 1st | 2nd | 2nd | 3rd | 5th | 15th | 7th | 2nd | 2nd | 4th | 9th | 8th | 16th | 12th | 14th |
| pH | 12th | 11th | 12th | 12th | 12th | 3rd | 3rd | 4th | 4th | 3rd | 17th | 18th | 19th | 18th | 18th |
| Rainfall 4-5 d | NA | NA | NA | NA | NA | 13th | 18th | 21st | 21st | 19th | 14th | 10th | 12th | 13th | 12th |
| Rainfall 0-1 d | NA | NA | NA | NA | NA | 19th | 16th | 16th | 15th | 7th | 20th | 20th | 20th | U | 19th |
| Rainfall 1-2 d | NA | NA | NA | NA | NA | 20th | 21st | 22nd | 20th | 22nd | 19th | 13th | 17th | 11th | 11th |
| Rainfall 2-3 d | NA | NA | NA | NA | NA | 9th | 8th | 8th | 9th | 8th | 11th | 9th | 4th | 6th | 10th |
| Rainfall 3-4 d | NA | NA | NA | NA | NA | 10th | 6th | 10th | 12th | 10th | 1st | 1st | 1st | 1st | 1st |
| Site | 14th | 12th | 10th | 10th | 15th | 2nd | 2nd | 3rd | 3rd | 2nd | 7th | 7th | 5th | 8th | 7th |
| Min. Since Midnight | 16th | U | U | 16th | 17th | 14th | 13th | 18th | 18th | 18th | NA | NA | NA | NA | NA |
| Turbidity | 8th | 10th | 9th | 9th | 9th | 6th | 5th | 6th | 6th | 6th | 16th | 16th | 15th | 16th | 13th |
| Water Temp. | 4th | 5th | 5th | 5th | 4th | 7th | 11th | 9th | 11th | 13th | 10th | 17th | 18th | 17th | 16th |
| Day of the Week | 17th | 16th | 16th | 17th | 16th | 5th | 4th | 5th | 5th | 5th | 2nd | 5th | 2nd | 9th | 4th |
| Year-day | 13th | 15th | 13th | 15th | 14th | 8th | 9th | 12th | 10th | 12th | 12th | 6th | 6th | 10th | 9th |
| Accuracy | 0.738 | 0.733 | 0.738 | 0.738 | 0.725 | 0.776 | 0.779 | 0.785 | 0.790 | 0.776 | 0.794 | 0.791 | 0.791 | 0.798 | 0.776 |
| Kappa Statistic | 0.473 | 0.464 | 0.474 | 0.474 | 0.449 | 0.482 | 0.497 | 0.511 | 0.516 | 0.487 | 0.238 | 0.239 | 0.164 | 0.193 | 0.177 |

Table S8: Results of random forests that identified factors associated with *E. coli* levels and likelihood of detecting pathogens in AZ and NY grab samples. Five, overlapping time frames [0-1, 0-2, 0-3, 0-4, or 0-5 d before sample collection (BSC)] were used to calculate the values of weather factors with the exception of rainfall; rainfall was calculated daily (0-1, 1-2, 2-3, 3-4, 4-5 d BSC). Separate random forests were developed for each outcome and overlapping time frame in each state.

| Factor | No. of Times in the Top 4 [a] | | | | | | | | Total Times in Top 4 [c] | | |
|---|---|---|---|---|---|---|---|---|---|---|---|
| | *E. coli* Levels | | *Salmonella* | | *eaeA-stx* Codetection | | *L. monocytogenes* | *Listeria* spp. [a] | | | |
| | AZ | NY | AZ | NY | AZ | NY | NY | NY | AZ | NY | Combined |
| pH | | 5 | | 3 | | 5 | 1 | 4 | 0 | 17 | 17 |
| Sample Site | 5 | | | | | 5 | | 5 | 5 | 10 | 15 |
| Sample Site | 5 | | | | | 5 | | 5 | 5 | 10 | 15 |
| Avg. Air Temperature | 5 | | 3 | 1 | 5 | | | | 13 | 1 | 14 |
| Max. Air Temperature | 1 | | 5 | 1 | 5 | | 1 | | 11 | 1 | 12 |
| Min. Air Temperature | 3 | 1 | | | 4 | 3 | | | 7 | 4 | 11 |
| Flow Rate | | 5 | | | | | 5 | 3 | 0 | 8 | 8 |
| Day of the Week | | | 5 | 5 | | 1 | 1 | | 5 | 5 | 6 |
| Dissolved Oxygen | 5 | 1 | | | | | | | 5 | 1 | 6 |
| Year-day | | | 2 | | | | 5 | 5 | 2 | 2 | 5 |
| Avg. Wind Speed | | 2 | | 2 | | | | 1 | 0 | 0 | 5 |
| Turbidity | | 5 | | | | | | | 0 | 5 | 5 |
| MPN of *E. coli*/100-mL | - [d] | - | | | | 5 | | | 0 | 5 | 5 |
| Rainfall 0-1 d BSC | - | | - | 4 | - | | | | 0 | 4 | 4 |
| Rainfall 4-5 d BSC | - | | - | 3 | - | | 1 | | 0 | 3 | 3 |
| Time of Day | 1 | | 2 | | | | | | 3 | 0 | 3 |
| Avg. Solar Radiation | | 1 | 1 | | 4 | 1 | 1 | | 8 | 5 | 2 |
| Avg. Relative Humidity | | | | 1 | | | 3 | 1 | 0 | 0 | 2 |
| Conductivity | | | 2 | | | | | | 2 | 0 | 2 |
| Water Temperature | | | | | 2 | | | 1 | 7 | 2 | 1 |
| Rainfall 3-4 d BSC | - | | - | | - | | 2 | | 0 | 0 | 0 |
| Rainfall 1-2 d BSC | - | | - | | - | | | | 0 | 0 | 0 |
| Rainfall 2-3 d BSC | - | | - | | - | | | | 0 | 0 | 0 |

[a] The number of random forests (out of 5) that the given factor was among the four top-ranked factors for the given outcome and state.
[b] Includes *L. monocytogenes.*
[c] The total number of times the given factor was among the four top-ranked factors across all outcomes and time-frames.

[d] - indicates that the given factor was not included as a covariate. For example, due to limited rainfall in AZ during the study period, rain was not included as a factor in the AZ RF. Therefore the total number of times a rainfall factor could have been ranked among the top four factors was 20.

Table S9: List of interactions between environmental factors that were investigated using two-way partial dependence plots. Due to the number of potential interactions that could have been investigated (e.g., 136 interactions could have been investigated for the random forests where the outcome was likelihood of *Salmonella* isolation in AZ grab samples), we focused on the impact of biologically plausible interactions on the likelihood of detecting pathogens in grab samples. For example, past studies have shown that the ability of solar radiation to damage bacterial cells is positively associated with dissolved oxygen levels (1–4).

| | Avg. Air Temp. | Dissolved Oxygen | *E. coli* Levels | Flow Rate | pH | Rainfall | Solar Radiation | Turbidity | Water Temp. |
|---|---|---|---|---|---|---|---|---|---|
| Avg. Air Temp. | | X | X | | | | | | |
| Dissolved Oxygen | | | X | | X | | X | | X |
| *E. coli* Levels | | | | X | X | X | X | X | X |
| Flow Rate | | | | | | | | X | |
| pH | | | | | | | | | |
| Rainfall | | | | | | | | X | |
| Solar Radiation | | | | | | | | X | |
| Turbidity | | | | | | | | | |
| Water Temp. | | | | | | | | | |
